# Dynamics of the habitat use of three predatory freshwater fish in a lentic ecosystem

**DOI:** 10.1101/2021.12.16.471647

**Authors:** Milan Říha, Ruben Rabaneda-Bueno, Ivan Jarić, Allan T. Souza, Lukáš Vejřík, Vladislav Draštík, Petr Blabolil, Michaela Holubová, Tomas Jůza, Karl Ø. Gjelland, Pavel Rychtecký, Zuzana Sajdlová, Luboš Kočvara, Michal Tušer, Marie Prchalová, Jaromír Seďa, Jiří Peterka

## Abstract

To understand the conditions of coexistence in multiple-species predator community, we studied longitudinal and vertical movement of pike (*Esox lucius*), pikeperch (*Sander lucioperca*) and catfish (*Silurus glanis*) in the Římov Reservoir, using an autonomous telemetry system for 11 months. We found significant differences among these three species in movement and depth that varied considerably in time, with the greatest differences between warm (late spring and early autumn) and cold season (late autumn to early spring). Preference for different sections of the reservoir was stable for pike, while pikeperch and catfish frequently visited tributary during the warm season, and moved closer to the dam during the cold season. Pike longitudinal activity was highest in the cold season, pikeperch in the warm season, and catfish activity peaked in both the warm and cold seasons. Overlap in the depth used among species was higher in the warm season, when all species used the upper layer of the water column, and lower in cold season, when pikeperch and catfish used deeper areas. These results demonstrated ability of predators to actively inspect a large portion of the reservoir in both longitudinal and vertical dimensions, although differing in the timing of their habitat use and activity.

## Introduction

Predators are an essential component of ecosystems because of their regulatory function (Carpenter, Kitchell, & Hodgson, 1985; Pelinson, Leibold, & Schiesari, 2021) and are often economically important species. This makes them highly relevant to a wide range of stakeholders, including ecosystem service managers, the scientific community and the general public. However, despite the great interest in freshwater predators and their ecology, there is relatively little information on their movement ecology and habitat use, as tracking their movements in large lentic waterbodies is challenging (Říha et al., 2021; Westrelin et al., 2021).

The spatial distribution of freshwater fish predators and their dynamics are determined by numerous abiotic and biotic factors, and individual and species characteristics and stage (Giske, Huse, & Fiksen, 1998). In general, predators should prefer and seek habitats with favorable hunting conditions and prey availability (Hugie & Dill, 1994). Such conditions vary among species according to their hunting strategies and adaptations and result in differences in predator activity and distribution (Pavlov & Kasumyan, 2002). These differences reduce competition among predators and ultimately lead to spatiotemporal habitat partitioning, with consequences for both predators and prey (Werner et al., 1977; Hughes & Grabowski, 2006).

Canyon-shaped reservoirs are complex systems with characteristic morphological and limnological features. They usually exhibit longitudinal gradients (from an inlet of an inflowing river to the dam) in various abiotic and biotic factors (Vašek et al., 2016). The most important ones are gradually increasing depth (from shallow, non-stratified tributary to deep, stratified sections near the dam) and decreasing nutrient concentration towards the dam. Such productivity gradients affect the entire system by producing similar gradients in primary producers (algae) and consumers (zooplankton and fish), as well as a wide range of environmental and foraging conditions, from highly trophic, turbid, and prey-rich sections near tributaries to lower trophic and more transparent sections with low prey abundance closer to the dam (Prchalová et al., 2009; Vašek et al., 2016). Thus, different predators with different hunting strategies may be favored in different longitudinal parts of the reservoir, and lead to a spatial segregation between such different predators.

The apex predators Northern pike (*Esox lucius*), pikeperch (*Sander lucioperca*), and wels catfish (*Silurus glanis*) are among the most abundant piscivorous predators in European freshwaters and have high ecological and economic value (Overton et al., 2015; Cucherousset et al., 2018; Skov & Nilsson, 2018). Previous studies have shown that the hunting conditions and movement patterns of these species differ considerably in several aspects of their ecology: (i) in the degree of site fidelity, with fidelity being highest for wels catfish and lowest for pikeperch (Fickling & Lee, 1985; Keskinen et al., 2005; Slavík et al., 2007; Brevé et al., 2014; Daněk et al., 2016; Sandlund, Museth, & Øistad, 2016; Cucherousset et al., 2018), (ii) in preferred hunting conditions, since Northern pike is a visually oriented predator (Skov & Nilsson, 2018), whereas pikeperch and wels catfish hunt at low light intensity (Cucherousset et al., 2018; Jokela-Määttä et al., 2019), and (iii) in temperature preference, with pikeperch and wels catfish being warm-water species, while Northern pike is considered a cool-water species (Feiner & Höök, 2015; Cucherousset et al., 2018; Skov & Nilsson, 2018). In addition, changes in seasonal activity and habitat preferences has been reported for all three species (Jepsen, Koed, & Økland, 1999; Slavík et al., 2007; Baktoft et al., 2012). Hence, when these three apex predators are living in sympatry in a reservoir, a spatial and temporal segregation driven by the longitudinal characteristics of the reservoir should be expected.

To understand if and how the longitudinal movement and depth use differ between Northern pike, pikeperch and wels catfish in sympatry, we used an autonomous telemetry system for 11 months in the Římov Reservoir (Czech Republic) to track the movements of tagged individuals of these species. We hypothesize that pikeperch and wels catfish spend more time in the high-turbidity area near the tributary than Northern pike. We expect that longitudinal activity will be highest for pikeperch and lowest for wels catfish. Furthermore, pikeperch and wels catfish should prefer shallower depths than Northern pike due to their preference for higher temperatures. Lastly, we assume that the differences among species in the parameters studied should vary temporally, with significant seasonal changes. In this study, we focus solely on the longitudinal, tributary-dam movement and depth use and neglect horizontal, inshore-offshore movement and diurnal aspect.

## Material and methods

### Study site

This study was conducted in deep, narrow, canyon-shaped the Římov Reservoir, frequently studied site (Znachor et al., 2016) with a single river inflow and stable abiotic conditions, as well as a well-developed longitudinal productivity gradient (Vašek et al., 2016). The reservoir is located in South Bohemia, Czech Republic (N 48°51.00978’, E 14°29.47462’; Fig. 1), constructed for drinking water storage and flood control. The reservoir was built in 1978 by damming the Malše River. The maximum area of the Římov Reservoir is 210 ha, the maximum volume is 33,106 m^3^ and the maximum depth is 45m with an average depth of 16 m. The length of the reservoir is about 8.5 km (measured along the central longitudinal axis of the reservoir) and the maximum surface altitude is 471 m a.s.l. The theoretical mean retention time is about 92 days. The reservoir is dimictic, with summer stratification usually lasting from April to September. There are no submerged aquatic macrophytes in the littoral zone due to steep banks and water level fluctuations. The trophic state of the reservoir is mesotrophic to eutrophic, with phosphorus and chlorophyll-a concentrations decreasing towards the dam (Seda & Devetter, 2000). Algal, zooplankton and fish density, and turbidity follow the trophic gradient, with the highest values near the tributary and decreasing towards the dam (Vašek et al., 2016).

**Figure 1.**
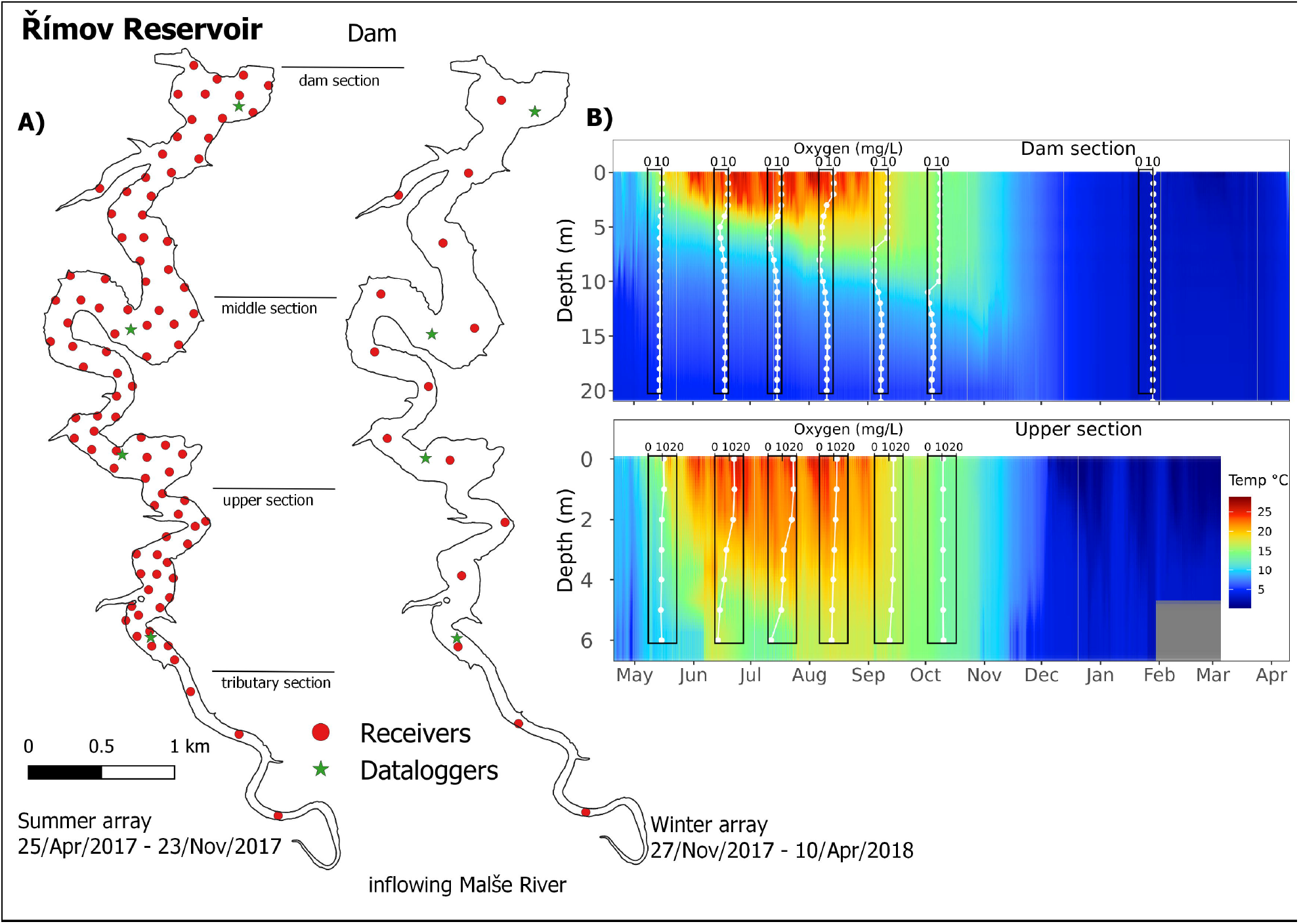
A) Positions of both telemetry arrays and dataloggers deployed in the Římov Reservoir. B) Development of temperature and oxygen stratification of water column at the dam and upper sites during the whole tracking period (note that the data loggers were deployed to a depth of 20 m at the dam site, but the maximum depth at this site is 42 m).

### Fish tagging

A total of 15 Northern pike and 20 pikeperch individuals were caught by electrofishing and 15 wels catfish individuals by long-lining. Northern pike were caught along the entire reservoir shore; pikeperch were caught at two locations in the reservoir and wels catfish were caught at four locations in the reservoir (Supplementary Material Fig. S1). Details of the fish capture are summarized in Table 1. After capture, all individuals were anesthetized with 2-phenoxy-ethanol (SIGMA Chemical Co., USA, 0.7 ml L^-1^, mean residence time in anesthetic bath 3.75 min), measured, weighed and tagged. A 1-1.5 cm incision was made on the ventral surface posterior to the pelvic girdle and a transmitter (Lotek Wireless Inc., MM-M-11-28-TP, 65×12 mm, mass in air 13 g, including pressure and temperature sensors, burst rate 15 s) was inserted through the incision and advanced into the body cavity. The incision was closed with two separate sutures. The mean surgery time was 3 min. All fish were released immediately after recovery from anesthesia at the site where they were captured. Fish were caught and tagged between April 18 and 25, 2017.

**Table 1.** Description of tagged fish in the Římov Reservoir. NTa stands for the number of tagged individuals, NAn for the number of analyzed individuals. Total length (TL), weight (W) and ratio of tag weight (TW) to fish body weight (FBW) are given for analyzed fish. Species NTa/NAn TL (cm)/W (kg) Ratio TW to FBW(%)

| Species | NTa/NAn | TL (cm)/W (kg) |  |  |  | Ratio TW to FBW(%) |  |  |
| --- | --- | --- | --- | --- | --- | --- | --- | --- |
|  |  | mean | SD | min | max | mean | min | max |
| pike | 15/10 | 70/2.9 | 17/2.8 | 50/0.8 | 116/10.9 | 0.5 | 0.1 | 1.7 |
| pikeperch | 20/7 | 49/1.0 | 6/0.4 | 40/0.5 | 56.5/1.6 | 1.3 | 0.8 | 2.5 |
| wels catfish | 15/14 | 117/12.3 | 26/9.5 | 86/3.8 | 166/35 | 0.1 | 0.03 | 0.3 |

### Fish tracking

MAP positioning system (Lotek Wireless Inc., Canada) was deployed in the reservoir to locate tagged fish. Two different arrangements of receiver arrays were consecutively deployed in the reservoir during the tracking period, to address the specific challenges of tracking fish in different seasons.

The first array was deployed in the reservoir from the start of tracking on 18 April 2017 to 20 November 2017 (hereafter referred to as the summer array). This array consisted of a total of 90 receivers (Lotek Wireless Inc., WHS3250) that were deployed to achieve complete coverage and fine scale positioning of the entire reservoir (86 receivers), except for the near tributary and the small bay near the dam (Fig. 1). Individual receivers were placed in the near tributary area (3 receivers) and in the small bay (1 receiver) to obtain presence data from these areas. The distance between receivers in the entire array was 80 - 300 m. The exact positions of the deployed receivers were measured using a differential GPS instrument, Spectra Precision, Promark 220 (USA). Based on range tests conducted prior to the reservoir survey (November 2016), this receiver array was arranged to provide fine-scale position data from the entire reservoir area, except the near tributary and the small bay monitored by individual receivers. Monitoring of systems accuracy was achieved using 22 stationary reference tags (Lotek Wireless Inc., Canada, model MM-M-16-50-TP, burst rate 25 sec) placed at 7 sites (in depths of 1, 5, and 10 m at all sites and in addition at 20 m near the dam). In addition, the accuracy of the entire system was tested after the system was deployed (July 2017) and before the final recovery of the system (November 2017) by dragging the reference tags across the reservoir by boat.

The second arrangement of the receiver array was deployed from 27 November 2017 to 10 April 2018 (referred to here as the winter array, Fig. 1). The winter array consisted of 15 receivers (of the same type as in the summer array) distributed along the centerline of the reservoir to track fish movement along the longitudinal axis of the reservoir at lower resolution. The reason for the lower resolution in winter was the inability to maintain the entire summer array when the reservoir was covered by ice. The performance and detection capability of the winter array was tested in December 2017 by dragging reference tags from a boat across the reservoir. Individual and total yield of fish locations during the study period is given in Table 2.

**Table 2.** Yield of positions (summer array) and detections (winter array) gathered by positioning systems during the whole tracking period for analyzed individuals (mean for all individuals, range among individuals and total sum of all gathered positions).

| Species | Positions (summer array) |  |  |  | Detections (winter array) |  |  |  |
| --- | --- | --- | --- | --- | --- | --- | --- | --- |
|  | mean | min | max | sum | mean | min | max | sum |
| pike | 133 527 | 60 437 | 185 702 | 1 335 270 | 105 032 | 49 420 | 148 436 | 840 258 |
| pikeperch | 138 481 | 110 097 | 194 001 | 969 371 | 115 401 | 92 923 | 139 593 | 577 009 |
| wels catfish | 142 228 | 54 969 | 195 669 | 1 991 205 | 119 459 | 52 654 | 183 029 | 1 552 978 |

### Temperature and oxygen monitoring

To obtain abiotic parameters that may influence the spatial distribution of the tracked species, we monitored water temperature and oxygen concentration in the reservoir. Water temperature was monitored using 60 data loggers (Onset, USA, HOBO Pendant temp/light 64K). The data loggers were placed at four locations to cover the longitudinal axis of the reservoir (Fig. 1). At each location, the data loggers were attached at 1 m intervals to a rope that extended from the surface to a depth of 13 m (data logger locations in dam and middle sections) or to 6 m (data logger in the upper section). An additional data logger was located at a depth of 20 m (data logger locations in dam and middle sections). The rope was attached to a floating buoy anchored to the bottom. This arrangement ensured both dense coverage at depths with rapid temperature changes and, with a 5 min measurement interval, high spatial and temporal resolution of the temperature profile. Oxygen concentration was measured at each data logger location each week during the summer array deployment and once during the winter array deployment (February 2018) using a calibrated YSI 556 MPS probe (YSI Incorporated, USA).

### Data processing

For the fine-scale summer array, individual horizontal fish positions were first calculated using the manufacturer’s proprietary positioning software UMAP v.1.4.3 (Lotek Wireless Inc., Canada). Fish depth was recorded from the tag’s internal sensor, which has a resolution of 0.7 m. In the next step, raw horizontal positions were filtered using a series of general additive models (GAM) and only positions that exceeded a threshold of 75 m beyond the final GAM were included in the analysis (details in Říha et al., 2021). The next step was to visually inspect the position estimates (unfiltered and filtered) and depth profiles of each fish. If both horizontal and vertical positions remained constant with no fish movement, it was interpreted as either a dead individual or a tag that was expelled, which happened for 5 Northern pike, 13 pikeperch, and 1 wels catfish, which were removed from further analyses.

In the final step, horizontal positions were projected onto the longitudinal center line of the reservoir by the shortest distance from the position to the line, and the longitudinal distance from the dam to the projected point along the center line was then calculated. For individual summer receivers (i.e., those near the tributary and a small bay), the position of each receiver was projected onto the centerline of the reservoir, and the distance from the dam along the centerline to these projected points were used as the longitudinal distance from the dam for individual fish detected by these receivers.

For the winter array, the distance of fish from the dam was calculated similarly to the individual receivers in summer. However, in some cases, the detection ranges of the receivers overlapped and duplicate detections occurred, i.e., an individual was detected by two or three receivers at the same moment. In such cases, the detection with the highest power value (detected in RSSI units, Received Signal Strength Indicator) was included in the analyses and the remaining detections were discarded, since we assumed that the fish was closest to the receiver receiving the strongest signal from the tag.

### Statistical analyses

Three movement parameters, preference for different parts of the reservoir, longitudinal activity and depth were calculated. The preferences for reservoir sections and their seasonal changes were discretized using the categories developed for the Římov Reservoir by Prchalová et al., (2009), with the reservoir divided into dam (0 – 1800 m from dam), middle (1800 - 5200 m), upper (5200 - 6600 m) and tributary area (6600 - 8650 m; Fig. 1; for more details about characteristics of each section see Prchalová et al., 2009). Longitudinal activity was defined as the difference between the maximum and minimum distance from the dam, for each individual and each day separately thus indicating daily range of longitudinal movement. Depth was calculated as the daily mean depth for each individual.

#### Reservoir section and depth use

Cumulative link mixed-effects models (CLMMs) were used to examine the effects of seasonality and temperature on reservoir section (CLMM_res)_ and depth (CLMM_depth_) use by the three species. CLMMs are a special class of general linear mixed-effects models that account for the effects of predictors on an ordered categorical outcome fitted with a logit link function (see SM for more details on CLMM parameterization). The response variables analyzed were: (1) reservoir section use (res_sec_use), categorised as dam, middle reservoir, upper reservoir, or tributary according to distance travelled by fish from 0-1800, 1800-5200, 5200-6600, or greater than 6600 m and coded as 1, 2, 3, and 4, respectively; and (2) depth use (depth_use), for which we considered residence at a given depth within a maximum daily range and was divided into seven levels corresponding to increasing water depths from 0-1 to over 10 m and coded as 1, 2, 3, 4, 5, 6, and 7, respectively. CLMMs were fitted using the *clmm* function of the R package *ordinal* (Christensen, 2019).

To estimate the differential effects of season and species on reservoir section use (res_sec_use) and depth (depth_use), we first used a likelihood ratio test to compare a model with the interaction between the categorical variables season (spring I April 27 – June 21 2017, summer June 21 – September 22 2017, autumn September 22 – December 21 2017, winter December 21 2017 - March 20 2018, and spring II March 20 2018 - April 10 2018) and species (Northern pike, pikeperch, and wels catfish) to a model without the interaction. If the season × species interaction was supported, we included it in the models. The fish identity tag (fishID) was included in the models as a random intercept to account for variability among fish.

The two CLMMs in this study can be expressed as follows:

CLMM_res_:

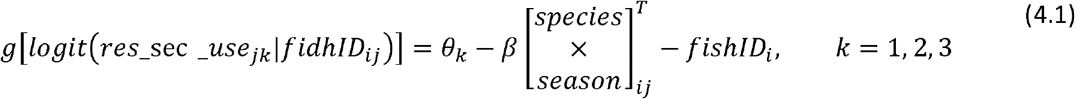

CLMM_depth_:

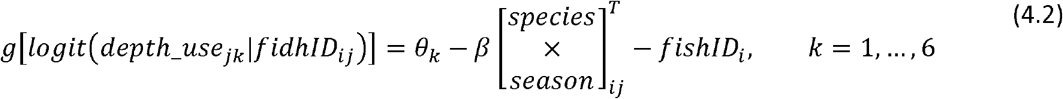

Here, θ_k_ is the cumulative logit for each response category *k*, which in this study takes the values *k* = 4 for the CLMMres representing four sections of the reservoir and *k = 7 for the* CLMM_depth_ representing increasing depth in the water column. fishID_i_ are subject-level random effects. β is a vector with three fixed effect estimates for each of the predictors species, season, and the season × species interaction. Therefore, β[species × season]ijT can be expressed as follows: β1(species) - β2(season) - β3(species × season) (see SM for more details on model definition).

#### Calculation of odds ratio from the fitted CLMMs

From the two fitted CLMMs, we calculated the odds ratio (OR) of Y_ij_ ≤ k, which is independent of the response category *k* as follows:

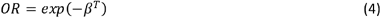

#### Longitudinal movement

We examined differences in longitudinal movements of Northern pike, pikeperch, and wels catfish from April 2017 to April 2018 and the effects of body length. Because our data span only one year and due to the lack of variability among years, we modelled the time series as a long-term trend component to increase the accuracy of the fish locations identified. Therefore, the time series included observations over the day of the year (with day 1 on 27 April 2017), while seasonality was graphically interpolated from the corresponding data. To trend the data for each species, we used a generalized additive mixed effects model (GAMM) with a smooth function for time and a random smooth function for fish tag identity as random=list(fishID =∼1) to account for variability among fish. These models were fitted separately as GAMM_pike_, GAMM_pikeperch_, and GAMM_catfish_. Although our time series does not include a seasonal component in a strict sense, we assume that fish move similarly in April 2017 and April 2018 due to seasonal preferences (e.g., temperature). Therefore, the values of the longitudinal range should be the same between the extremes of the time series. Moreover, the cyclical property allows us to estimate the rate of change of the curve using the continuous domain of the cubic spline and its first derivatives. To account for the cyclic assumption, we added a cubic regression spline (with the argument bs = “cc”) to the time spline function. To control for possible effects of body length (“bl”) on longitudinal motion, we fitted a model that also included a smoothing function for this covariate and compared it with a likelihood ratio test to a model without this covariate.

The equation of this model can be expressed as follows:

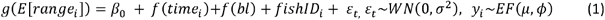

Where *g* is an identity link function, *E[range_i_]* is the expected value of the response variable with mean *μ* and variance of the scale parameter *⍰, β_0_* is the estimated model intercept, *⍰*_*i*_ is the random effect for *fishID*_*i*_, *f* are smoothing functions for the trend and bl covariates, and *ε*_*i*_ is a random error term of the residual temporal correlation, where *W* is a covariance matrix with mean 0 and variance *σ*^*2*^ (see SM for more details on the GAMM parameterization).

#### Identification of periods with significant changes

We performed a functional data analysis to examine how the longitudinal range changed over time. To identify periods with significant upward and downward trends, i.e., the rate of change (slope) of the nonlinear time trend, the periods with statistically significant changes were determined using the finite difference method. In this approach, the value of the fitted spline function for the trend component is determined by calculating the first derivative of a time *t*, i.e., the slope between each two closely spaced time points, given by:

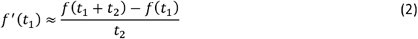

The first derivative gives an idea of the change in response relative to the change between these two adjacent time points, which would represent the expected differences in longitudinal trends (see SM for more details on the first derivative calculations). Given this premise, we used the *predict*.*gam* function and the argument type = “lpmatrix” to generate a set of prediction data from the fitted GAMM to evaluate the change in the function. We then used the *derivatives* function of the *gratia* package (Simpson, 2020) to calculate the first derivatives of the estimated smoothing values.

We calculated the simultaneous confidence intervals using the Bayesian covariance matrix with smoothing correction (Simpson, 2020). In this trend analysis, there is a significant change, either up or down, in a given period of the time series if the estimated confidence intervals do not include the zero value in that period. To obtain better precision in estimating the derivative, we evaluated the slope with an interval of about 3 hours between each pair of time points. Considering that our data contained observations every minute, this is a precise value while ensuring reasonable computation times.

#### Smooth differences between species

We fitted a new GAMM (GAMM_sp_diffs_) to examine whether the three species exhibited different longitudinal movements away from the dam at different times of the year. In this model, we used the full data set and added the factor variable “species” for the time-spline function with the argument “by” (s(time, by = species)). Including this smoothing factor interaction (time × species) estimates a separate smoothing for each species with its own smoothing parameter (λ_i_) and penalty term, which we can use to estimate the difference between the fitted trends. Since each estimated smoothing function is subject to identifiability constraints, with a separate penalty that shrinks it in the direction of the null effect, to identify periods with significant differences in longitudinal movement we also included the factor variable “species” as a parametric term in the model. This was also used to estimate the mean of the longitudinal range for each species. Expanding Eq. (1) the formula of this model is expressed as:

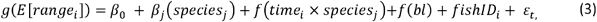

Where β_j_ is the mean intercept value of the response for a given level of species_j_.

We used the plot_diff() function from the package “itsadug” (van Rij et al., 2017) to determine the differences between each pair of the three estimated smoothings. This function evaluates the smoothings over the entire time domain at each species by generating a set of predicted values that are used to evaluate the basis functions of the model and plot the difference curves. The uncertainty of the estimated difference is calculated using the variance-covariance matrix of the coefficient values estimated from the model as the standard error of the estimated pairwise difference. We calculated simultaneous confidence intervals based on Bayesian posterior simulations of the fitted GAMM, where the covariance matrix of the smoothing parameters includes a correction for the uncertainty in their estimates (with the argument unconditional= TRUE).

#### Residual autocorrelation in fitted GAMMs

Since in our daily time series the residuals of successive time points are likely to be highly interdependent, we applied an ARMA process to the residuals in the formula GAMM to handle these serial correlations. We chose an autoregressive AR (2) term after comparing it to two models, one with AR (1) and one without autocorrelation using a generalized likelihood ratio test. The model AR (2) also performed better in modelling temporal correlation based on the normalized residuals. The autoregressive term was included in the model formula with the correlation argument as corARMA(form = ∼ 1|time, p = 2).

All models were fitted using restricted maximum likelihood estimation (REML) with Gaussian error distribution and identity link function using the function gamm() from the package “mgcv” (Mixed GAM Computation Vehicle with Automatic Smoothness Estimation, Wood, 2021) implemented in R software version 3.6.3 (R Core Team, 2020).

## Results

### Reservoir sections use

Considering the average use of reservoir sections over time for the three species, there were significant changes between seasons, except between late spring and summer (LSM across seasons ± SE: −0.05 ± 0.09, z = −0.58, *p* = 0.98) and between autumn and winter (LSM: 0.08 ± 0.08, z= 0.88, *p* = 0.90; Table 3 and S3 in SM). The pattern of reservoir section use was similar for pikeperch and wels catfish (Fig. 2), and both species preferred the dam and middle sections in autumn, winter, and early spring. However, as predicted by the model, the upper section and tributary were used more in late spring (P(Y_j_ ≥ 3) ± SE: upper reservoir – pikeperch: 0.41 ± 0.01, wels catfish: 0.40 ± 0.06; tributary – pikeperch: 0.28 ± 0.13, wels catfish: 0.40 ± 0.17) and in summer (upper reservoir – pikeperch: 0.40 ± 0.04, wels catfish: 0.37 ± 0.08; tributary – pikeperch: 0.23 ± 0.11, wels catfish: 0.46 ± 0.18), with no significant trend in change between the two seasons for either pikeperch or wels catfish (LSM of seasonal differences: both, *p* > 0.1). Northern pike used sections of the reservoir similarly throughout the tracking period, except in early spring (Fig. 2). From late spring through winter, Northern pike preferred the middle and upper sections, while in early spring they preferred the dam and middle sections. Thus, the probability of using these reservoir sections increased significantly from winter onward (LSM across season × species: 2.09 ± 0.34, z = 6.14, *p* < 0.001) and decreased toward late spring (LSM: −1.99 ± 0.34, z = 5.83, *p* < 0.001) (Figs. 2 and S2, S3 in SM).

**Table 3.** Summary of the cumulative link mixed model predicting differences in reservoir sections use by pike, pikeperch and wels catfish across seasons from April 2017 to April 2018. Numbers represent odds ratios (OR) and 95% confidence intervals (95% CI). res_sec is an ordinal dependent variable used to fit the CLMM_res_ model, with *k* = 4 categories related to different reservoir sections, where θ_k_ is the cumulative probability of using each category (dam, middle reservoir, upper reservoir and tributary). *β*_1_*-*_3_ are exponentials of the estimated fixed effects regression coefficients for each of the interaction covariates species × season (see main text and Supplementary Information for a description of the variables and model fit). Random effects are represented by the individual intercepts for the fish identity tag (fishID) (τ_00_) and the residual variance (σ^2^). ICC is the intraclass correlation coefficient, which measures the degree of repeatability at the individual level. Marginal R^2^/Cond. R^2^ are marginal and conditional r-squared values that refer to the proportion of variation explained by fixed effects and the variance explained by fixed and random effects, respectively. In bold, *p* < 0.05.

| Variable | OR [95% CI] | z | p |
| --- | --- | --- | --- |
| **res_sec (thresholds)** |  |  |  |
| $\theta_1$ (dam \| middle) | 0.12 [0.01 – 1.29] | - | 0 |
| $\theta_2$ (middle \| upper) | 1.23 [0.12 – 12.89] | 0 | 0 |
| $\theta_3$ (upper \| tributary) | 7.19 [0.69 – 75.34] | 1 | 0 |
| **Species** |  |  |  |
| $\beta_1$ (pike) (ref.) | | - | - |
| $\beta_1$ (pikeperch) | 2.83 [0.20 – 40.63] | 0 | 0 |
| $\beta_1$ (wels catfish) | 4.74 [0.30 – 75.39] | 1 | 0 |
| **Season** |  |  |  |
| $\beta_2$ (spring I) (ref.) | | - | - |
| $\beta_2$ (summer) | 1.23 [0.84 – 1.78] | 1 | 0 |
| $\beta_2$ (autumn) | 1.9 [1.31 – 2.74] | 3 | **0** |
| $\beta_2$ (winter) | 1.1 [0.74 – 1.63] | 0 | 0 |
| $\beta_2$ (spring II) | 0.14 [0.07 – 0.27] | - | < |
| **Species × Season** |  |  |  |
| $\beta_3$ (Northern pike × spring I) (ref.) | | - | - |
| $\beta_3$ (pikeperch × summer) | 0.61 [0.39 – 0.94] | - | **0** |
| $\beta_3$ (wels catfish × summer) | 1.04 [0.66 – 1.64] | 0 | 0 |
| $\beta_3$ (pikeperch × autumn) | 0.1 [0.06 – 0.16] | - | < |
| $\beta_3$ (wels catfish × autumn) | 0.09 [0.06 – 0.14] | - | < |
| $\beta_3$ (pikeperch × winter) | 0.24 [0.14 – 0.38] | - | < |
| $\beta_3$ (wels catfish × winter) | 0.15 [0.09 – 0.24] | - | < |
| $\beta_3$ (pikeperch × spring II) | 1.99 [0.86 – 4.59] | 1 | 0 |
| $\beta_3$ (wels catfish × spring II) | 1.56 [0.71 – 3.42] | 1 | 0 |
| **Random Effects** |  |  |  |
| $\sigma^2$ | 3.29 | | |
| $\tau_{00}$ fishID | 2.53 | | |
| ICC | 0.43 |  |  |
| $N_{\text{fishID}}$ | 13 | | |
| Marginal $R^2/\text{Cond. } R^2$ | 0.104 / 0.493 | | |

**Figure 2.**
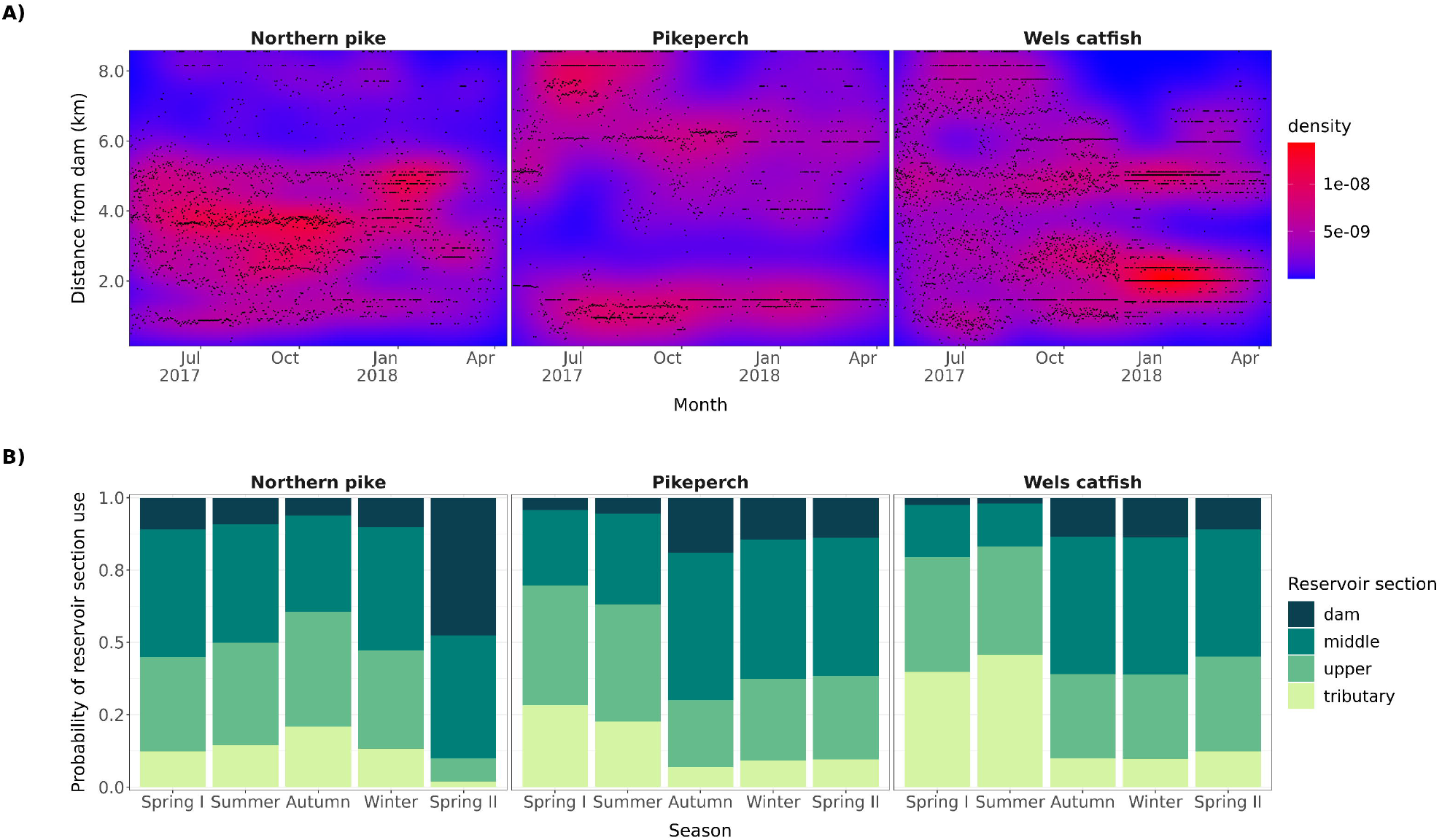
**(A)** Density of locations of all tagged individuals in relation to distance from dam and time **(B)** Cumulative probabilities of reservoir section use in Římov Reservoir by Northern pike, pikeperch and wels catfish at different seasons of the year from April 2017 to April 2018, estimated using a cumulative link mixed-effects model analysis (CLMM_res_). The y-axis represents the probability of fish using each section of the reservoir at different times of the year (x-axis). The dam represents the starting point of the measured longitudinal migration distance and is set as the reference category in the analysis.

### Longitudinal movement

The range of the total explored reservoir area throughout the tracking period did not differ significantly among species (Fig. 3). Movement of most individuals of all three species ranged from 50 to 75 % of the extent of the reservoir (4.2 to 6.4 km). Intraspecific differences were greater than interspecific differences and slightly higher for Northern pike (15 – 91 %; 1.3 - 7.8 km) and wels catfish (17 – 91 %; 1.5 - 7.8 km) than for pikeperch (37 – 94 %; 3.2 - 8 km; Fig. 3).

**Figure 3.**
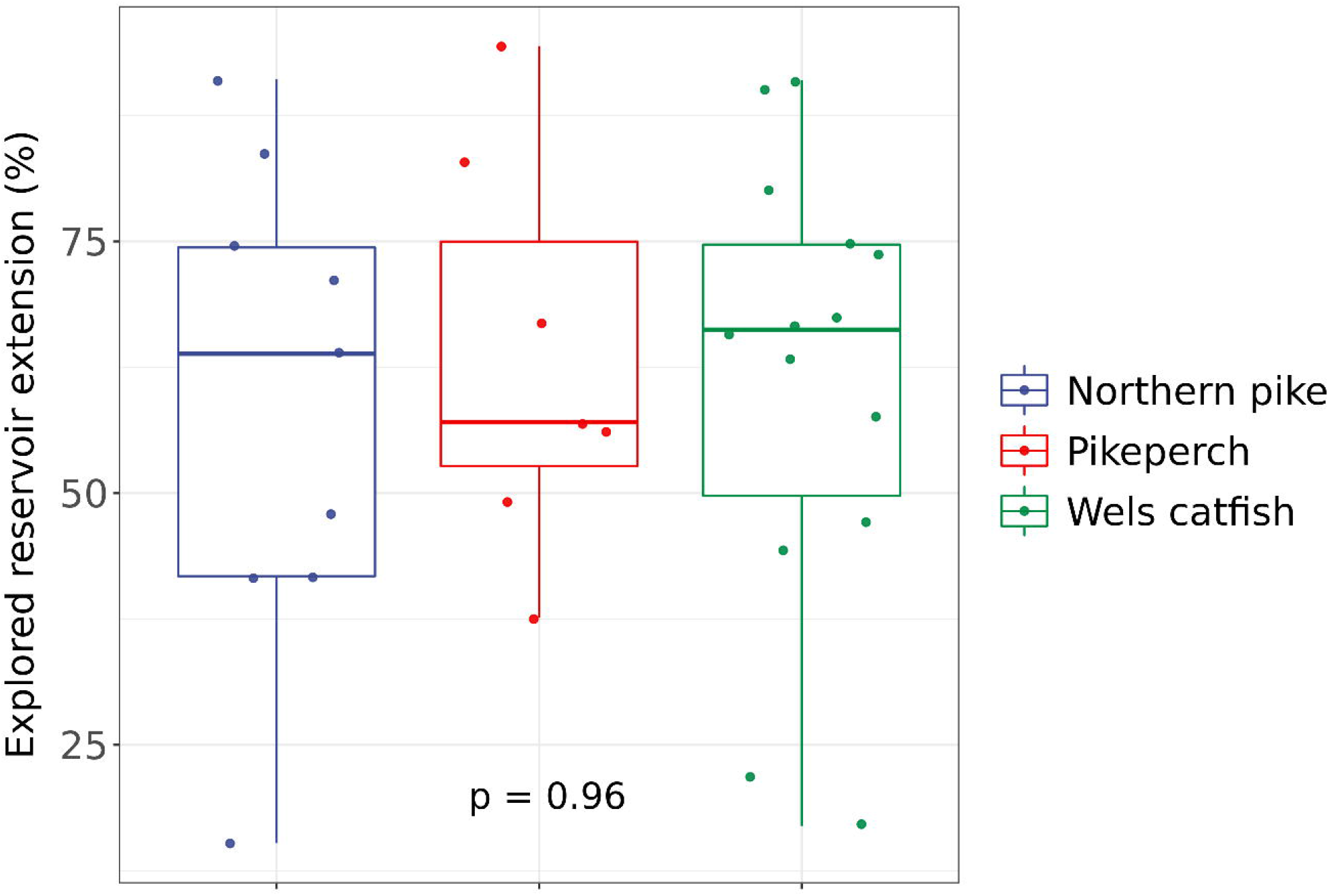
Explored length of the Římov reservoir during the whole tracking period (April 2017 to April 2018) by each individual (dots) and significance of ANOVA test comparing interspecific differences in this parameter.

Smooth functions of long-term trends were significant in all fitted models (all, p < 0.001) and the time-spline functions show variability among species over time, suggesting that the temporal pattern of longitudinal activity differs among species (Tab. 4, 5; Fig. 4a). Wels catfish covered the greatest average longitudinal distance per day (mean ± SE: 0.99 ± 0.09 km/day, t=10.55, *p* < 0.001), followed by Northern pike (0.90 ± 0.11 km/day, t=8.045, *p* < 0.001) and finally pikeperch (0.77 ± 0.13 km/day, t=5.75, *p* < 0.001).

**Table 4.** Results of generalized additive mixed model analysis of differences in longitudinal movements of pike, pikeperch, and wels catfish from April 2017 to April 2018. The model (GAMMsp_diffs) was fitted with a separate smoothing trend for each species by including a factor smoothing interaction term in the time-spline function (s(Time)). To avoid limitations in model identifiability, species was also included as a parametric term in the analysis. Fish identity identifier (fishID) was included as random smooth intercepts to account for variability among fish (see Statistical analysis for more details on model fit). R^2^ adj, adjusted r-squared. ⍰ is the estimated scale parameter associated with the variance of the random effects. In bold, p < 0.05.

| **A. parametric coefficients** | **Estimate** | **Std. Error** | **t-value** | **p-value** |
| --- | --- | --- | --- | --- |
| (Intercept) | 0.90 | 0.11 | 8.04 | **<0.001** |
| pikeperch | -0.13 | 0.17 | -0.75 | 0.453 |
| wels catfish | 0.10 | 0.15 | 0.65 | 0.514 |
| **B. smooth terms** | **edf** | **Ref.df** | **F-value** | **p-value** |
| s(Time) × pike | 7.20 | 8.00 | 92.95 | **<0.001** |
| s(Time) × pikeperch | 5.74 | 8.00 | 116.86 | **<0.001** |
| s(Time) × wels catfish | 7.61 | 8.00 | 42.47 | **<0.001** |
| s(fishID) | 27.44 | 28.00 | 52.55 | **<0.001** |
| **$R^2$ adj** | 0.21 |  |  |  |
| **Deviance explained (%)** | 21 |  |  |  |
| **Scale parameter ($\sigma^2$)** | 0.64 |  |  |  |

**Table 5.**
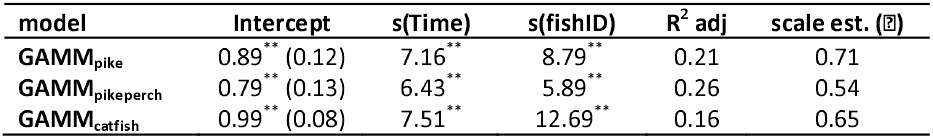
GAMM analysis of longitudinal data on pike, pikeperch, and wels catfish movements from April 2017 to April 2018. Models (GAMMpike, GAMMpikeperch, GAMMcatfish) were fitted separately for each species dataset using a cubic regression spline for the smooth function of time trend (s(Time)) and a random smooth function for fish tag identity (s(fishID)) (see Statistical Analysis for more details on model fitting). Numbers for the parametric components (intercept) refer to estimates (standard errors). The numbers for the smoothing components refer to the estimated effective degrees of freedom (edf), which reflect the degree of nonlinearity/complexity of the relationship between a covariate and the response. ⍰ is the estimated scale parameter estimated from a GAMM associated with the variance of the random effects. R^2^ adj, adjusted r-squared. * p < 0.05; ** p < 0.01.

**Figure 4.**
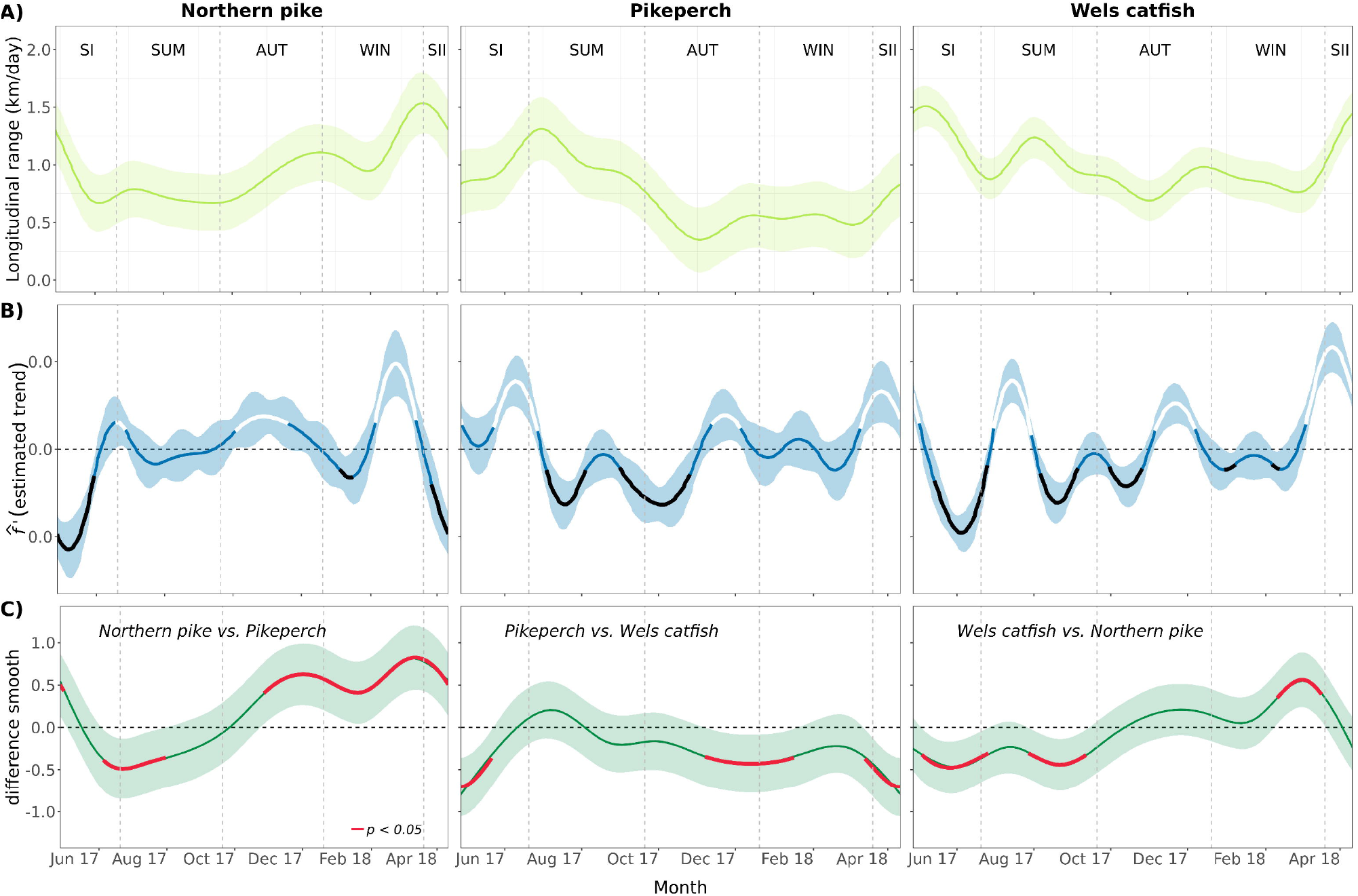
**A)** Effect of smooth time functions on longitudinal movement (km.day^-1^) of pike, pikeperch and wels catfish in Římov Reservoir in different seasons. The fitted spline represents the average trend effect from April 2017 to April 2018 based on the additive mixed-effects models GAMM_pike_, GAMM_pikeperch_, and GAMM_catfish_ with AR (2) residual error term. Body length was not included as a covariate in any of the models because it had insignificant effects on longitudinal movement (see Results for more details). The shaded area represents the 95% CI for the fitted penalised spline function. **(B)** Rate of change in longitudinal movements over time, determined from the estimated first derivative of the fitted trend spline function from each GAMM. The periods of statistically significant (zero is not included in the 95% confidence interval) increasing (thick white) and decreasing (thick black) rates of change in the trend series are depicted. The shaded area is the 95% simultaneous confidence interval calculated using posterior Bayesian simulations for the data predicted from the models. **C)** Estimation of smooth differences in longitudinal movement trends between each species. Shaded area indicates 95% confidence intervals. Periods where the value zero (horizontal line) is not included in the 95% confidence interval indicate significant differences between two species (outlined by a red line). Differences in s(time) by (species) interactions between each species pair are significant at *p* < 0.05.

The first-derivative analysis of the rate of change in longitudinal movements over time identified several periods of significant (decreasing or increasing) change (Fig. 4b). The longitudinal activity of Northern pike decreased significantly in late spring (April 27–May 30), increased slightly at the beginning of summer (June 18–27), and increased again significantly in autumn (October 3–November 19) before decreasing for almost two weeks (January 4–16). This was followed by an increase from mid-winter to early spring (February 5–March 13) before declining significantly (March 27–April 11; Fig. 4b). It is worth noting that the increase was much steeper in winter than in autumn, peaking around February 25.

Pikeperch activity increased significantly in late spring and early summer (May 23–June 27), peaked in July, and then declined markedly and steadily during a month in summer (July 4–August 4) and during an extended period from late summer through mid-autumn (August 31–November 26). Pikeperch activity increased significantly during mid-autumn (November 12–December 5), but remained consistently low during winter and early spring, and increased again during a month from late winter to early spring (March 10–April 11; Fig. 4b).

Wels catfish activity showed a significant downward trend in late spring through the beginning of summer (May 14–June 25), followed by a significant increase in July (June 30–July 30). Activity then declined significantly, bottoming out in two consecutive periods (August 6–September 5 and September 30–October 26). Activity resumed in early November (November 5–December 7), followed by two short periods of about a week with significantly lower activity (December 30–January 8 and February 6– February 12), and finally followed by a period of significantly increased activity from late winter to late spring (March 5–April 11), peaking in April (Fig. 4b).

Trend analysis of smooth differences among species (GAMM_sp_diffs_) revealed exact periods of significant differences in their longitudinal movements from the dam (Fig. 4c). Significant differences were found between Northern pike and pikeperch from mid-autumn (October 27) to late spring (April 11) and from the end of late spring (June 4) to mid-summer (July 31) (difference smooth: 5.70, F = 116.34, *p* < 0.001). Differences between Northern pike and wels catfish were in late spring (May 4) to the beginning of summer (June 25), from mid-summer (July 27) to the end of summer (September 11) and from mid-winter (February 10) to the end of winter (March 17) (difference smooth: 7.60, F = 41.77, *p* < 0.001). Pikeperch and wels catfish differed from the beginning of late spring to mid-late spring (April 27 to May 21), from the beginning of spring 2018 (March 13 to April 11), and from mid-autumn 2017 (November 6) to mid-winter 2018 (January 16) (difference smooth: 7.20, F = 92.95, *p* < 0.001).

### Depth use

When considering the average use of depth over time for the three species, no significant differences were found in late spring (spring I) and summer (least-squares means, LSM, of pairwise comparisons across season × species: all, *p* > 0.1), i.e., when they preferred the epilimnetic part of the water column up to 5 m (Table 6 and S6 in SM, Fig. 5). However, within this range, the near-surface area (depth to 2 m) was used more in late spring than in summer (LSM across seasons ± SE: −1.6 ± 0.09, z = −18.90, *p* < 0.001), as predicted by the model (P(Y_j_ ≤ 2) ± SE: spring I: 0.42 ± 0.03 vs. summer: 0.28 ± 0.04) and preference for the shallowest waters (down to 1 m depth) decreased significantly from late spring to summer for all species (OR: 12.4 [8.50 – 18.08], *p* < 0.001). A similar pattern to that observed in late spring was observed in early spring for Northern pike and pikeperch, while wels catfish showed a gradually increasing probability of using greater depths down to 10 m in early spring (OR: 90.63 [40.59 – 202.32], *p* < 0.001), with 7-10 m being the preferred depth range (P(Y_j_ ≤ 6) ± SE: 0.35 ± 0.08; Fig. 5).

**Table 6.**
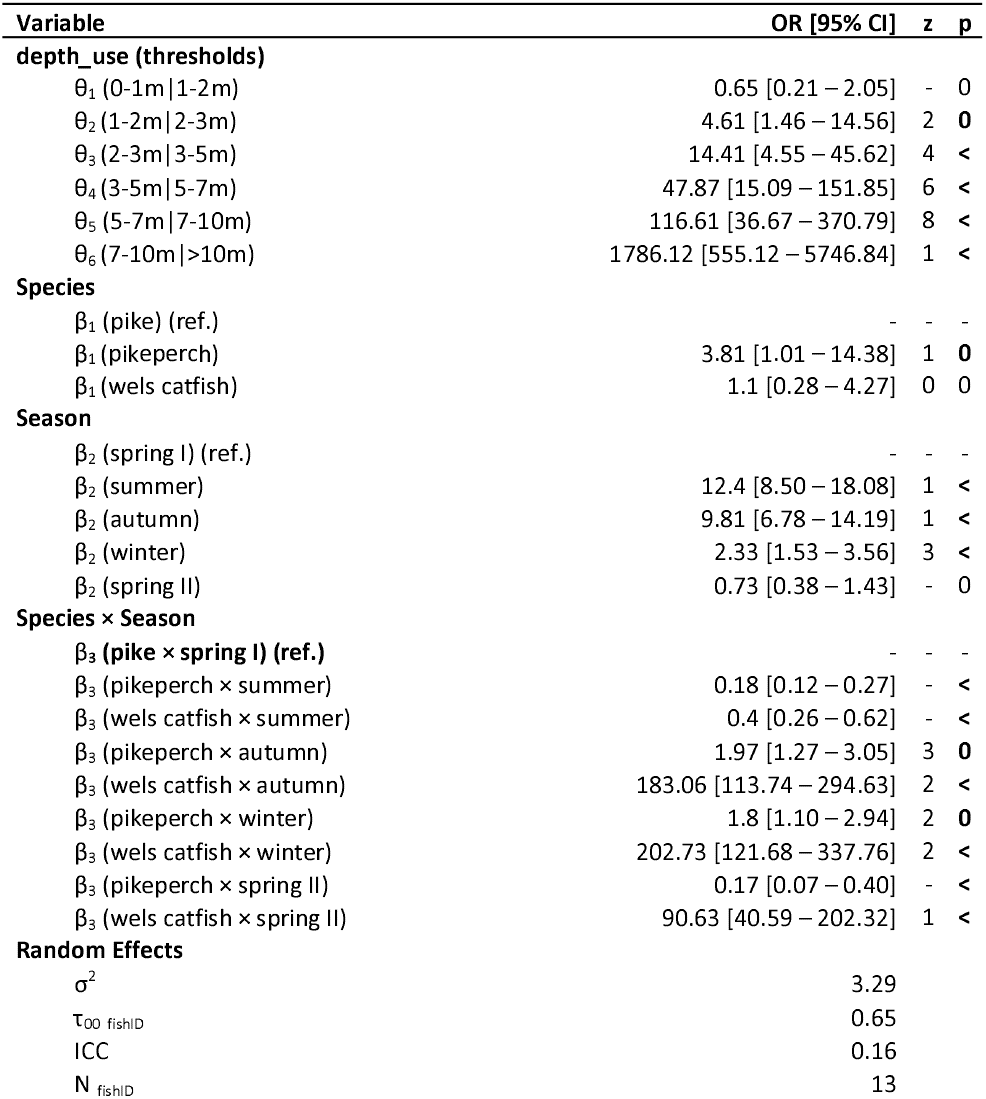
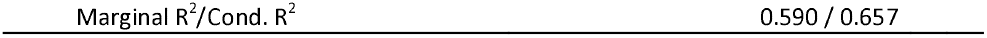
Summary of the cumulative link mixed model predicting differences in depth use by Northern pike, pikeperch and wels catfish across seasons from April 2017 to April 2018. Numbers represent odds ratios (OR) and 95% confidence intervals (95% CI). depth_use is an ordinal dependent variable used to fit the CLMM_depth_ model, with *k* = 7 categories related to different depth ranges, where θ_k_ is the cumulative probability of using each category (from 0-1 to > 10 m). *β*_1_*-*_3_ are exponentials of the estimated fixed effects regression coefficients for each of the interaction covariates species × season (see main text and Supplementary Information for a description of the variables and model fit). Random effects are represented by the individual intercepts for the fish identity tag (fishID) (τ_∞_) and the residual variance (σ^2^). ICC is the intraclass correlation coefficient, which measures the degree of repeatability at the individual level. Marginal R^2^/Cond. R^2^ are marginal and conditional r-squared values that refer to the proportion of variation explained by fixed effects and the variance explained by fixed and random effects, respectively. In bold, *p* < 0.05.

**Figure 5.**
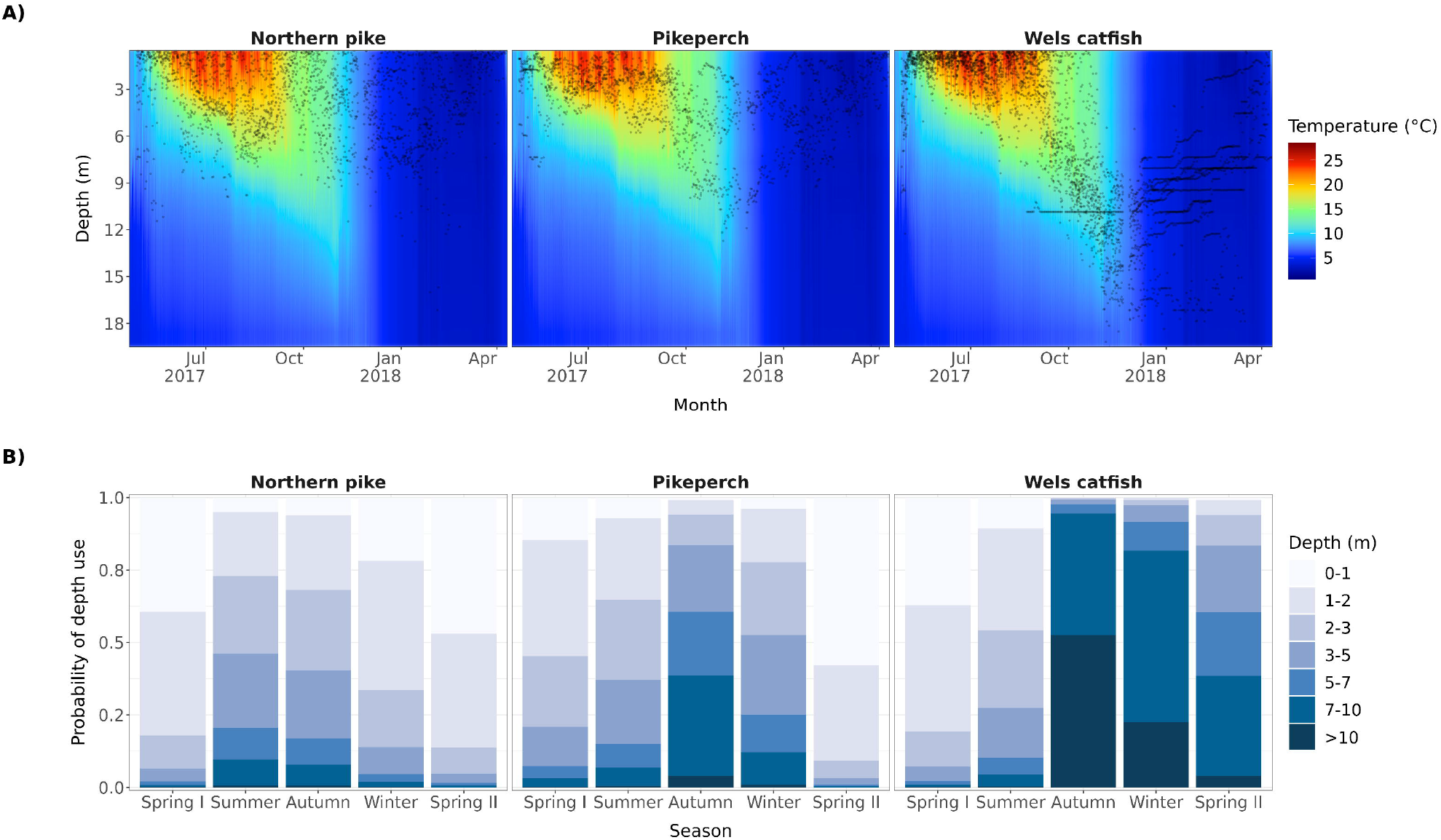
**(A)** Detected mean daily depths of all tracked individuals (black dots) in relation to temperature (colours) and time; **(B)** Cumulative probabilities of depth use in Římov Reservoir by Northern pike, pikeperch and wels catfish at different times of the year from April 2017 to April 2018, estimated with a cumulative link mixed-effects model analysis (CLMM_depth_). Depth was categorized into seven levels corresponding to increasing depth in the water column, from 0-1 to > 10 m. The y-axis represents the probability of using the depth at each season (x-axis). Depth 0-1 m was set as the reference category in the analysis.

During the destratification of the water column (early autumn), species differed markedly in their depth use. In autumn, Northern pike used a similar depth as in summer (mean depth 3 m) (LSM across seasons ± SE: 0.23 ± 0.16, z = 1.48, *p* < 0.98), and both varied slightly but significantly compared to winter (summer-winter: 1.67 ± 0.20, z = 8.47, *p* < 0.001; autumn-winter: 1.44 ± 0.19, z = 7.44, *p* < 0.001). Pikeperch and wels catfish used deeper water in autumn than in winter, with the former more likely to use depths of 5 to 10 m in autumn (OR: 1.97 [1.27 - 3.05], *p* = 0.002) and 3 to 5 m in winter (OR: 1.8 [1.10 - 2.94], *p* = 0.018), while the second were significantly more likely to use depths greater than 10 m in autumn (OR: 183.06 [113.74 - 294.63], *p* < 0.001) and 7 to 10 m depth in winter (OR: 202.73 [121.68 – 337.76], *p* < 0.001) in winter (Figs. 5 and S4 SM).

## Discussion

Our study revealed significant differences in habitat use among the three predatory species – Northern pike, pikeperch and wels catfish, in terms of their preference for different sections, longitudinal activity and use of depth and in an 8.5 km long reservoir. Interspecific differences in these parameters varied considerably over time, with the greatest differences between the warm season (late spring and early autumn) and the cold season (late autumn to early spring). Preference for different sections of the reservoir was stable for Northern pike, while pikeperch and wels catfish frequently visited tributary and the upper sections of the reservoir during the warm season and moved closer to the dam during cold season. Overall longitudinal activity was similar for all species, but Northern pike activity was highest in the cold season, pikeperch activity peaked in the warm season, and wels catfish activity had several peaks in both the warm and cold seasons. Overlap in depth use among species was greatest in warm season, when all species used the upper layers of the water column, and least in cold season, when pikeperch and especially wels catfish used deeper areas.

### Preference for particular reservoir sections

Our results showed that the preference for reservoir sections was dependent on species and season. The strongest changes in pikeperch and wels catfish were found in the sections closer to the inflow of Malše River (upper and tributary sections). These two species frequently visited these reservoir sections in late spring and summer, and then gradually moved closer to the dam as the stratification of the water column disappeared. The upper and especially tributary sections of the reservoir were more strongly avoided in cold season by wels catfish than by pikeperch. Wels catfish gradually moved closer to the dam through January and then gradually retreated toward the tributary for the rest of the winter and early spring. For pikeperch, the section near the tributary was completely avoided only in October and November, after which some individuals occasionally revisited this section. For both species, the timing of these changes corresponded to changes in their depth preference (as described above). Overwintering in deeper areas closer to the dam has been previously documented for pikeperch (Jepsen, Koed, & Økland, 1999), but to our best knowledge, such behavior has not been documented for wels catfish in lentic environments.

Previous studies in the Římov Reservoir described a stable longitudinal gradient in productivity closely associated with turbidity and prey fish distribution during warm season (Prchalová et al., 2009; Vašek et al., 2016). These gradients make the sections near the tributary of the Malše River rich in prey fish, but also eutrophic and hence turbid. Pikeperch and wels catfish are species well adapted to turbid conditions (Cucherousset et al., 2018; Jokela-Määttä et al., 2019), so their preference for these sections may be related to the presence of favorable conditions and higher prey density. Their avoidance of this area during the cold season may be related to their preference for greater depths during that period, which are not available in the shallow upper sections and the tributary. However, other factors could also be responsible for these behavioral differences, such as seasonal changes in prey density or other intra- and interspecific interactions, which need to be further investigated.

Northern pike seemed to avoid the upper section of the reservoir. Moreover, the preference of reservoir sections was stable across seasons, changing only in early spring when they moved nearer to the dam. The timing of changes in Northern pike preference for particular sections of the reservoir corresponds to Northern pike spawning activity (Pauwels et al., 2014) and it is very likely that these changes in preference are related to the location of suitable spawning habitat.

### Longitudinal activity

Our 11-month tracking showed that the upper reach of the reservoir was much less used by pike then by wels catfish and pikeperch, although the longitudinal extent visited was generally similar for all species, covering 50-75% (4-6.5 km) of the reservoir extent. These results are consistent with previous studies on pikeperch (Fickling & Lee, 1985; Koed, 2001; Vehanen & Lahti, 2003), but show greater movement ability for Northern pike and wels catfish than is usually documented (Cucherousset et al., 2018), and corroborate with recent studies that showed greater space use and movement by these species (Capra, Pella, & Ovidio, 2018; Nyqvist et al., 2020; Lenhardt et al., 2021; Říha et al., 2021). It shows that these species were able to survey a relatively large portion of the reservoir and select suitable locations to reside during different parts of the annual cycle, and partly question the view of Northern pike and wels catfish as stationary species with a relatively strict home range (Slavík et al., 2007; Craig, 2008; Brevé et al., 2014; Daněk et al., 2016; Sandlund, Museth, & Øistad, 2016), while pikeperch is a species with low site fidelity (Fickling & Lee, 1985; Koed, 2001; Vehanen & Lahti, 2003).

Despite the similarities in overall longitudinal coverage, we found large differences among species in the temporal pattern of longitudinal activity. Telemetry studies on Northern pike reported ambiguous differences in warm and cold seasons movement activity. They showed similar or higher winter activity (Jepsen et al., 2001; Koed et al., 2006; Baktoft et al., 2012; Nordahl et al., 2020) with a peak in activity in early spring (Pauwels et al., 2014) or reduced activity in the cold season (Cook & Bergersen, 1988; Rogers & Bergersen, 1995; Kobler et al., 2009). Our results support studies showing higher winter activity, as Northern pike longitudinal activity was low in summer, gradually increased in the cold season, and peaked in early spring.

Pikeperch longitudinal activity peaked in summer (July and August) and was relatively low in other seasons, in accordance with other findings from lentic waters (Jepsen, Koed, & Økland, 1999; Vehanen & Lahti, 2003), and in contrast to observation of riverine pikeperch with highest activity in spring and autumn (Koed, 2001; Horký, Slavík, & Bartoš, 2008). We expected a similar temporal pattern for wels catfish, since it has been previously documented that they have their highest activity in spring and summer and are inactive (or even hibernate) during the cold season (Daněk et al., 2014, 2016; Cucherousset et al., 2018; Kuzishchin, Gruzdeva, & Pavlov, 2018). However, our results showed that activity of wels catfish was similar throughout the tracking period, with several peaks in late spring, summer, and the transition between autumn and winter. These findings corroborate the results of recent studies suggesting lower but persistent wels catfish activity even during the cold season (Lenhardt et al., 2021; Santis & Volta, 2021).

Seasonal changes in activity are related to different phases of the annual life cycle, such as foraging, spawning, or overwintering, and their transitions (Horký et al., 2006; Slavík et al., 2007; Baktoft et al., 2012). The transition between foraging and overwintering may explain the differences between warm and cold seasons and the changes in pikeperch activity. Previous studies have found that pikeperch stop foraging at lower temperatures. Pikeperch have been found to feed significantly less below 8°C and barely at all below 4°C (Malinovskyi et al., 2019), although Teletchea et al., (2009) has shown that feeding rates during the cold season may depend on body fat content. A similar reduction in activity between foraging and hibernation was expected in wels catfish (Cucherousset et al., 2018). Wels catfish have been documented to stop feeding at temperatures below 7-10°C in aquaculture (David, 2006; Copp et al., 2009), or have been suspected to do so based on stomach studies (Wysujack & Mehner, 2002) and their reduced catches (Britton et al., 2007). However, recent study of Santis & Volta (2021) documented feeding even during winter, at temperatures below 10°C, and our results showed that wels catfish were active longitudinally in winter and that these movements were related to depth use (see below) and preference for particular reservoir sections. These changes in winter activity cannot be explained by changes in abiotic conditions (i.e, temperature, oxygen concentrations, or lack of currents), as they were homogeneous in the reservoir during winter. A more likely explanation is that foraging or other biotic interactions (with other conspecifics or other predators; Cucherousset et al., 2018) drove winter activity of wels catfish. However, winter ecology of freshwater fishes is generally poorly understood, and very little information is available on the winter movements and habitat preferences of species and their prey (Marsden et al., 2021).

In Northern pike, a reduced but sustained food intake has been documented even in winter (Diana, 1979), and it has been suggested that this is related to energy requirements for ovarian recruitment and early spring spawning (Baktoft et al., 2012). Based on these results, we can hypothesize that the increased movement of Northern pike along the reservoir during the cold seasons may be related to foraging. The highest peak in longitudinal activity of Northern pike was recorded in early spring in the Římov Reservoir. Such a pattern has been documented previously (Pauwels et al., 2014) and is thought to be caused by spawning activity, as the species spawns in early spring (Skov & Nilsson, 2018).

### Depth use

Our results showed a large overlap in depth use among species in spring and summer and their separation in depth use in autumn and winter. During the warm season, all three species used the upper part of the water column, which agrees well with the extent of the thermocline and confirms previous results (Copp et al., 2009; Nordahl et al., 2020; Říha et al., 2021; Westrelin et al., 2021). There could be several explanations for a similar depth preference in summer, which are not mutually exclusive and which may work together. The strong temperature stratification of the water column meant that the optimal temperatures of all three species (Feiner & Höök, 2015; Cucherousset et al., 2018; Skov & Nilsson, 2018) occurred only in the surface epilimnetic layer (down to 5 - 7 m during the period from late spring to early autumn). Moreover, anoxic conditions prevailed in most of the reservoir below the thermocline from late August, making the depths below the thermocline inaccessible to fish. In addition to abiotic factors, prey distribution could also play a crucial role in the distribution of aquatic predators (Brodersen, Howeth, & Post, 2015; Říha et al., 2021). Previous studies of the Římov Reservoir have shown nearly all prey species to be restricted to the epilimnion during the summer (Prchalová et al., 2009; Vašek et al., 2016). Therefore, the reason for the high overlap of depth use among predators in summer and early autumn was probably a consequence of the combined effects of temperature profile, oxygen availability and prey distribution.

Pikeperch and wels catfish responded similarly to the autumnal mixing of the water column, preferring greater depths after the thermocline diminished. Pikeperch used the deepest parts of the reservoir only in the autumn, while wels catfish gradually descended and reached the greatest depths in January. The preference for greater depths during the cold season for overwintering has been previously documented for these two species (Lehtonen, 1983; Lehtonen & Toivonen, 1988; Nyberg, Degerman, & Sers, 1996; Jepsen, Koed, & Økland, 1999; Daněk et al., 2014; Kuzishchin, Gruzdeva, & Pavlov, 2018). However, previous studies generally documented static behaviour for both species when overwintering in deep holes and assumed that this was a response to oxygen availability or currents (Lehtonen, 1983; Lehtonen & Toivonen, 1988; Nyberg, Degerman, & Sers, 1996; Jepsen, Koed, & Økland, 1999; Daněk et al., 2014; Cucherousset et al., 2018; Kuzishchin, Gruzdeva, & Pavlov, 2018). Such explanations cannot be applied to the Římov Reservoir (no current, homogeneous oxygen concentration), and we can hypothesize that preference for deeper areas may be a species-specific behavioural trait that is independent of conditions, at least in wels catfish. In pikeperch, not all individuals descended to greater depths, and some remained at the same depth during the cold period. However, other factors such as changes in prey distribution might play a role for both pikeperch and wels catfish, as preference of deeper river holes was documented for various species in riverine environment (Rakowitz et al., 2014). In the absence of a vertical temperature gradient (which has been suggested as a major factor in summer vertical distribution), the gradual ascent of wels catfish to shallower depths during late winter and early spring must be explained by other factors, such as the photoperiod duration or changes in the prey distribution.

### Caveats

Our study has several potential limitations that must be considered when interpreting the results. Sample size and body size range were limited for all three species. Therefore, statistical power to detect differences among species or more detailed analyses of reservoir section preferences were limited. We did not examine fish sex. However, sex-specific differences in activity have been documented previously, particularly during spawning in all three species (Jepsen, Koed, & Økland, 1999; Poulet et al., 2005; Copp et al., 2009; Pauwels et al., 2014). We did not include sex in the analyses, and it could therefore represent another factor contributing to the observed high interindividual variability. Differences in accuracy of summer and winter arrays could have led to a slight overestimation of winter activity compared to summer on individual species level. However, the winter array setup was the same for all three species, and the results for winter should therefore well reflect interspecific differences. Moreover, the seasonal changes in longitudinal activity of pikeperch and wels catfish agree well with the timing of their vertical movement (detected by internal sensors on tags, which were independent of the array deployment method) and confirm our conclusions about changes of their activity.

### Conclusions

Simultaneous automatic tracking of three predator species provided an opportunity to compare the timing and extent of their activities and the overlap in their space use. The study supported our hypothesis that pikeperch and wels catfish prefer the area near tributaries more than Northern pike. Our second hypothesis that longitudinal activity would be highest for pikeperch and lowest for wels catfish was not confirmed, as activity was similar for all species. Also, our hypothesis of depth differentiation with pikeperch and wels catfish preferring shallower depths than Northern pike was not confirmed, as all species preferred the same depths during temperature stratification of water column. Space use in the reservoir and habitat preferences may have a large influence on interactions among these predators, as well as between the predators and their prey. The three predator species studied have a large overlap in their prey in reservoirs with similar prey compositions (Vejřík et al., 2017; Adámek et al., 2019). Considering their spatial overlap, our results showed that they share similar spatial niche in the Římov Reservoir in spring and summer, using only the epilimnion. Moreover, pikeperch and wels catfish used more intensively the narrow tributary section, which could lead to their higher competition in this area. During the cold season (autumn and winter), the spatial overlap was reduced as they preferred different depths. Our study also showed that all three predators were also active in winter, dynamically changing their longitudinal and vertical locations. As cold season abiotic conditions were rather stable in the reservoir, such activity suggests foraging or other biotic interactions, which can be important for population dynamics and interspecies interactions (McMeans et al., 2020). However, there are critical gaps in our knowledge of winter ecology as there is no information how predator activity is related to their prey activity.

## Acknowledgements

Authors would like to acknowledge all FishEcU members for their help during fieldwork and they would like to thank to Vilém Děd for his help with data processing. The work was supported from ERDF/ESF project Biomanipulation as a tool for improving water quality of dam reservoirs (No. CZ.02.1.01/0.0/0.0/16_025/0007417) and the project QK1920011 “Methodology of predatory fish quantification in drinking-water reservoirs to optimize the management of aquatic ecosystems”. IJ acknowledges the sponsorship provided by the J. E. Purkyně Fellowship of the Czech Academy of Sciences.

## Statements and declarations

### Author’s contributions

All authors contributed substantial comments during manuscript preparation:

Conceptualization: MŘ

Data curation: MŘ

Formal analysis: RRB, MŘ, KØG

Statistical analysis: RRB

Funding acquisition: MŘ, JP, PB

Investigation: MŘ, LV, IJ, ATS, LV, V.Draštík, PB, MH, TJ, PR, ZS, LK, MT, MP, MS, JP

Methodology: MŘ, KØG, IJ, AST

Visualization: RRB, MŘ

Writing – original draft: MŘ, RRB

Writing – review & editing: all authors

### Availability of supporting data and materials

The datasets used and analyzed during the current study are available from the corresponding author on reasonable request.

