## Supplementary materials for "Dynamics of the habitat use of three predatory freshwater fish in a lentic ecosystem"

**SUPLEMENTARY MATERIALS**

**Extended Methods**

Data analyses

1. Longitudinal movement
   1.1. Model definition

We modelled the time series as a daily trend component rather than using a numerical monthly variable. Using a monthly trend would reduce accuracy because there would be no variability between years to model a seasonal component. Modelling it as a long-term trend would lose accuracy compared to using a daily series. Therefore, we only modelled a long-term change by using day-of-year observations.

A general model formula can be expressed as follows:

$$g\left( E\left[ y_{i} \right] \right)=\beta_{0} + f\left( x_{i} \right)+\gamma_{i}+ \varepsilon_{t},y_{i}\sim EF\left( \mu,\phi\right)$$

(1)

Where *g()* is an identity link function, *E*(y_i_) is the expected value of the response variable *y_i_,* for cluster *i* at time t_i_, assumed to be conditionally distributed with mean *μ* and variance equivalent to the scale parameter *ϕ; β_0_* is the average estimated effect (model intercept); *f_1_* is a smoothing function for the covariate X_1i_ in cluster *i*; *ϒ_i_* is the subject-level random factor with an i.i.d. Gaussian random coefficient; *ε_i_* is a random error term of the residual temporal correlation of the time series where *W* is a covariance matrix for a 2^nd^ order autocorrelation coefficient with mean 0 and variance σ^2^.

In this study, *ϒ_i_* corresponds to the random effect for fishID_i_ (intercept) with variable number of individuals for each species, *f_1_* are passed to the *s()* terms as penalized regression spline functions for the trend time_1i_ and body length covariates bl_1i_, and *ε_i_* is an ARMA process of order 2. By substituting Equation 1 the GAMM model has the following form:

$$g\left( E\left[ {long\_range}_{i} \right] \right)=\beta_{0} + s\left( {time}_{i} \right){+s\left( {bl}_{i} \right)+fishID}_{i}+ \varepsilon_{t,} \varepsilon_{t}\sim WN\left( 0,\sigma^{2} \right)$$

(2)

1.2. Identifying periods of significant change

For the smooth time function f(x), its rate of change between two time points t_1_ and t_2_ is given by the ratio between the change in the longitudinal values in that interval and the corresponding change in the time series values. Therefore, the derivative at time t_1_ is the instantaneous rate of change of f(x) as the time interval between the two time points decreases (i.e., t_2_ approaches 0), given by:

$$f^{'}\left( t_{1} \right)=\lim_{t_{2}\to0} \left( \frac{f\left( t_{1}+t_{2} \right)-f\left( t_{1} \right)}{t_{2}} \right)$$

(3.1)

$$f^{'}\left( t_{1} \right)\approx\frac{f\left( t_{1}+t_{2} \right)-f\left( t_{1} \right)}{t_{2}}$$

(3.2)

1.3. Smooth differences between species

We hypothesised that fish of the three species moved away from the dam at different times during the 1-year period and thus exhibited different longitudinal movements. To compare species pairs we used plot_diff() that is used to assess smooth differences. Based on the fitted GAMM, the predicted values are stored in the form of an Xβ matrix containing the estimated covariance matrices of both the coefficient values and the fitted model values over the time series (see R code).

2. Reservoir section and depth use

2.1. Definition of cumulative link mixed models

CLMMs are fitted with a logit link function *g()* given by:

$$g\left( logit\left[ \pi\right] \right)=log\left[ \frac{\pi}{1-\pi} \right]$$

(4)

For the ordinal response *Y* each j_th_ observation takes a value for the entire range of ordered categories *k* = {1,..., k_n_} and the probability that the j_th_ observation falls into each of the response categories is denoted by *π_jk_* as:

$$y_{jk}=P\left( Y_{j}\leq k \right)=\frac{P\left( Y_{j}\leq k \right)}{1-P\left( Y_{j}\leq k \right)}=\pi_{j1}+\ldots+ \pi_{jk}$$

(5)

Where P(*Y*_j_ ≤ *k*) is the cumulative probability of *Y* and P(*Y*_j_ ≤ *k*) /1- P(*Y*_j_ ≤ *k*) is the probability of being less than or equal to a given category *k*. Based on Equation 5, the cumulative logits of *Y* are the log-odds of being less than or equal to a given category *k* and they exclude the last category (i.e. k_n_ -1) *:*

$$logit\left( y_{jk} \right)=logit\left( P\left( Y_{j}\leq k \right) \right)= log\frac{P\left( Y_{j}\leq k \right)}{1-P\left( Y_{j}\leq k \right)}, k=1,\ldots., k_{n-1}$$

(6)

2.2. Parameterization of CLMM_res_ and CLMM_depth_

Using Equation 6, we extend the model to our data by including the fish identity tag (fishID) as a random intercept to account for inter-individual variability:

$$g\left( logit\left[ P\left( Y_{ij}\leq k|\mu_{i} \right) \right] \right)=\theta_{k}-\chi_{ij}^{T}\beta-\omega_{ij}^{T}\mu_{j}, k=1,\ldots., k_{n-1}, \mu_{j}\sim N\left( 0,\sigma^{2} \right)$$

(7)

Where *θ_k_* is the cumulative logit for each response category *k* with separate intercepts *θ_k_* = {θ_1k_ ,..., θ_nk_}, *χ_ij_* is a vector of fixed effects of the covariates for the *j_th_* observation in a given *fishID_i_* (i.e., where each fish individual i assumes a variable number for each species), *β* is a vector of regression coefficients, *μ_j_* is a vector of subject-level random effects for fishID_i_ that are i.i.d. Gaussian with mean E[*u_j_*]= 0 and variance *σ_u_^2^*, and *ω_ij_* is the corresponding set of random effects scaling parameters for the j_th_ observation in fishID_i_. k_n_ refers to the number of response categories, which in this study take the values k_n_ = 4 for the CLMM_res_ representing four sections of the reservoir and k_n_ = 7 for the CLMM_depth_ representing increasing depth in the water column.

Summarizing Equation 7, the two CLMMs in this study are parameterized as follows:

CLMM_res_:

$$g\left[ logit\left( {res\_sec\_use}_{jk}|{fidhID}_{ij} \right) \right]=\theta_{k}-\beta\left[ \begin{matrix} species \\ \times\\ season \end{matrix} \right]_{ij}^{T}-{fishID}_{i}, k=1, 2, 3$$

(8.1)

CLMM_depth_:

$$g\left[ logit\left( {depth\_use}_{jk}|{fidhID}_{ij} \right) \right]=\theta_{k}-\beta\left[ \begin{matrix} species \\ \times\\ season \end{matrix} \right]_{ij}^{T}-{fishID}_{i}, k=1, \ldots, 6$$

(8.2)

In addition to the CLMM_res_ and CLMM_depth_ models we also fitted a model with species and temperature as continuous variables without random effects using the clm function in the same package to predict changes in reservoir sections according to the water temperature gradient.

2.3. Calculation of odds ratio from the fitted CLMMs

From the two fitted CLMMs, we calculated the odds ratio (OR) of *Y*_ij_ ≤ *k*, which is independent of the response category *k*, as follows:

$$OR=\frac{exp\left( \theta_{k}-\chi_{1}^{T}\beta\right)}{exp\left( \theta_{k}-\chi_{2}^{T}\beta\right)}$$

(9)

Since the cumulative OR between levels {χ_1_,χ_2_ ,..., χ_n_} of a categorical variable *χ^T^* (e.g., OR of *Y*_ij_ ≤ *k* in season[spring I]_ij_) relative to *Y*_ij_ ≤ *k* in season[summer]_ij_) is proportional to the distance between the two levels, then χ_1_-χ_2_=1 ,..., χ_n_-χ_n-1_=1 and the OR is given by:

$$OR=exp\left( -\beta^{T} \right)$$

(10)

**Extended Results**

**Extended Table S3.** Summary of the cumulative link mixed model predicting differences in reservoir sections use by pike, pikeperch and wels catfish across seasons from April 2017 to April 2018. Numbers represent log odds (Estimate) and standard errors (S.E.). res_sec is an ordinal dependent variable used to fit the CLMM_res_ model, with *k* = 4 categories related to different reservoir sections, where θ_k_ is the cumulative probability of using each category (dam, middle reservoir, upper reservoir and tributary). *β_1_-_3_* are ordered log odds estimates of the fixed effects for each of the Species × Season interaction covariates (see main text and Supplementary Information for a description of the variables and model fit). Random effects are represented by the individual intercepts for the fish identity tag (fishID) (τ_00_) and the residual variance (σ^2^). ICC is the intraclass correlation coefficient, which measures the degree of repeatability at the individual level. Marginal R^2^/Cond. R^2^ are marginal and conditional r-squared values that refer to the proportion of variation explained by fixed effects and the variance explained by fixed and random effects, respectively. In bold, *p* < 0.05.

| Variable | Estimate | S.E. | z*-*value | *p-*value |
| --- | --- | --- | --- | --- |
| res_sec (thresholds) |  |  |  |  |
| θ_1_ (dam\|middle) | -2.09 | 1.2 | -1.74 | 0.081 |
| θ_2_ (middle\|upper) | 0.21 | 1.2 | 0.17 | 0.863 |
| θ_3_ (upper\|tributary) | 1.97 | 1.2 | 1.65 | 0.1 |
| Species |  |  |  |  |
| β_1_ (pike) (ref.) | - | - | - | - |
| β_1_ (pikeperch) | 1.04 | 1.36 | 0.77 | 0.443 |
| β_1_ (wels catfish) | 1.56 | 1.41 | 1.1 | 0.27 |
| Season |  |  |  |  |
| β_2_ (spring I) (ref.) | - | - | - | - |
| β_2_ (summer) | 0.2 | 0.19 | 1.07 | 0.284 |
| β_2_ (autumn) | 0.64 | 0.19 | 3.41 | **0.001** |
| β_2_ (winter) | 0.1 | 0.2 | 0.48 | 0.634 |
| β_2_ (spring II) | -1.99 | 0.34 | -5.83 | **<0.001** |
| Species × Season |  |  |  |  |
| β_3_ (pike × spring I) (ref.) | - | - | - | - |
| β_3_ (pikeperch × summer) | -0.5 | 0.23 | -2.22 | **0.027** |
| β_3_ (wels catfish × summer) | 0.04 | 0.23 | 0.17 | 0.864 |
| β_3_ (pikeperch × autumn) | -2.32 | 0.23 | -9.92 | **<0.001** |
| β_3_ (wels catfish × autumn) | -2.44 | 0.23 | -10.65 | **<0.001** |
| β_3_ (pikeperch × winter) | -1.45 | 0.25 | -5.86 | **<0.001** |
| β_3_ (wels catfish × winter) | -1.9 | 0.24 | -7.86 | **<0.001** |
| β_3_ (pikeperch × spring II) | 0.69 | 0.43 | 1.6 | 0.109 |
| β_3_ (wels catfish × spring II) | 0.44 | 0.4 | 1.11 | 0.266 |
| Random Effects |  |  |  |  |
| σ^2^ | 3.29 |  |  |  |
| τ_00_ _fishID_ | 2.53 |  |  |  |
| ICC | 0.43 |  |  |  |
| N _fishID_ | 13 |  |  |  |
| Marginal R^2^/Cond. R^2^ | 0.10/0.49 |  |  |  |

**Extended Table S6.** Summary of the cumulative link mixed model predicting differences in depth use by pike, pikeperch and wels catfish across seasons from April 2017 to April 2018. Numbers represent log odds (Estimate) and standard errors (S.E.). depth_use is an ordinal dependent variable used to fit the CLMM_depth_ model, with *k* = 7 categories related to different depth ranges, where θ_k_ is the cumulative probability of using each category (from 0-1 to > 10 m). *β_1_-_3_* are ordered log odds estimates of the fixed effects for each of the Species × Season interaction covariates (see main text and Supplementary Information for a description of the variables and model fit). Random effects are represented by the individual intercepts for the fish identity tag (fishID) (τ_00_) and the residual variance (σ^2^). ICC is the intraclass correlation coefficient, which measures the degree of repeatability at the individual level. Marginal R^2^/Cond. R^2^ are marginal and conditional r-squared values that refer to the proportion of variation explained by fixed effects and the variance explained by fixed and random effects, respectively. In bold, *p* < 0.05.

| Variable | Estimate | S.E. | z-value | *p*-value |
| --- | --- | --- | --- | --- |
| depth_use (thresholds) |  |  |  |  |
| θ_1_ (0-1m\|1-2m) | -0.43 | 0.59 | -0.74 | 0.462 |
| θ_2_ (1-2m\|2-3m) | 1.53 | 0.59 | 2.6 | **0.009** |
| θ_3_ (2-3m\|3-5m) | 2.67 | 0.59 | 4.54 | **<0.001** |
| θ_4_ (3-5m\|5-7m) | 3.87 | 0.59 | 6.57 | **<0.001** |
| θ_5_ (5-7m\|7-10m) | 4.76 | 0.59 | 8.06 | **<0.001** |
| θ_6_ (7-10m\|>10m) | 7.49 | 0.6 | 12.56 | **<0.001** |
| Species |  |  |  |  |
| β_1_ (pike) (ref.) | - | - | - | - |
| β_1_ (pikeperch) | 1.34 | 0.68 | 1.97 | **0.049** |
| β_1_ (wels catfish) | 0.1 | 0.69 | 0.14 | 0.89 |
| Season |  |  |  |  |
| β_2_ (spring I) (ref.) | - | - | - | - |
| β_2_ (summer) | 2.52 | 0.19 | 13.09 | **<0.001** |
| β_2_ (autumn) | 2.28 | 0.19 | 12.11 | **<0.001** |
| β_2_ (winter) | 0.85 | 0.22 | 3.93 | **<0.001** |
| β_2_ (spring II) | -0.31 | 0.34 | -0.91 | 0.361 |
| Species × Season |  |  |  |  |
| β_3_ (pike × spring I) (ref.) | - | - | - | - |
| β_3_ (pikeperch × summer) | -1.72 | 0.22 | -7.95 | **<0.001** |
| β_3_ (wels catfish × summer) | -0.92 | 0.23 | -4.02 | **<0.001** |
| β_3_ (pikeperch × autumn) | 0.68 | 0.22 | 3.04 | **0.002** |
| β_3_ (wels catfish × autumn) | 5.21 | 0.24 | 21.46 | **<0.001** |
| β_3_ (pikeperch × winter) | 0.59 | 0.25 | 2.36 | **0.018** |
| β_3_ (wels catfish × winter) | 5.31 | 0.26 | 20.4 | **<0.001** |
| β_3_ (pikeperch × spring II) | -1.78 | 0.43 | -4.08 | **<0.001** |
| β_3_ (wels catfish × spring II) | 4.51 | 0.41 | 11 | **<0.001** |
| Random Effects |  |  |  |  |
| σ^2^ | 3.29 |  |  |  |
| τ_00_ _fishID_ | 0.65 |  |  |  |
| ICC | 0.16 |  |  |  |
| N _fishID_ | 13 |  |  |  |
| Marginal R^2^/Cond. R^2^ | 0.590 / 0.657 |  |  |  |

**Extended figures**

**Figure S1.** Captures locations of tagged individuals

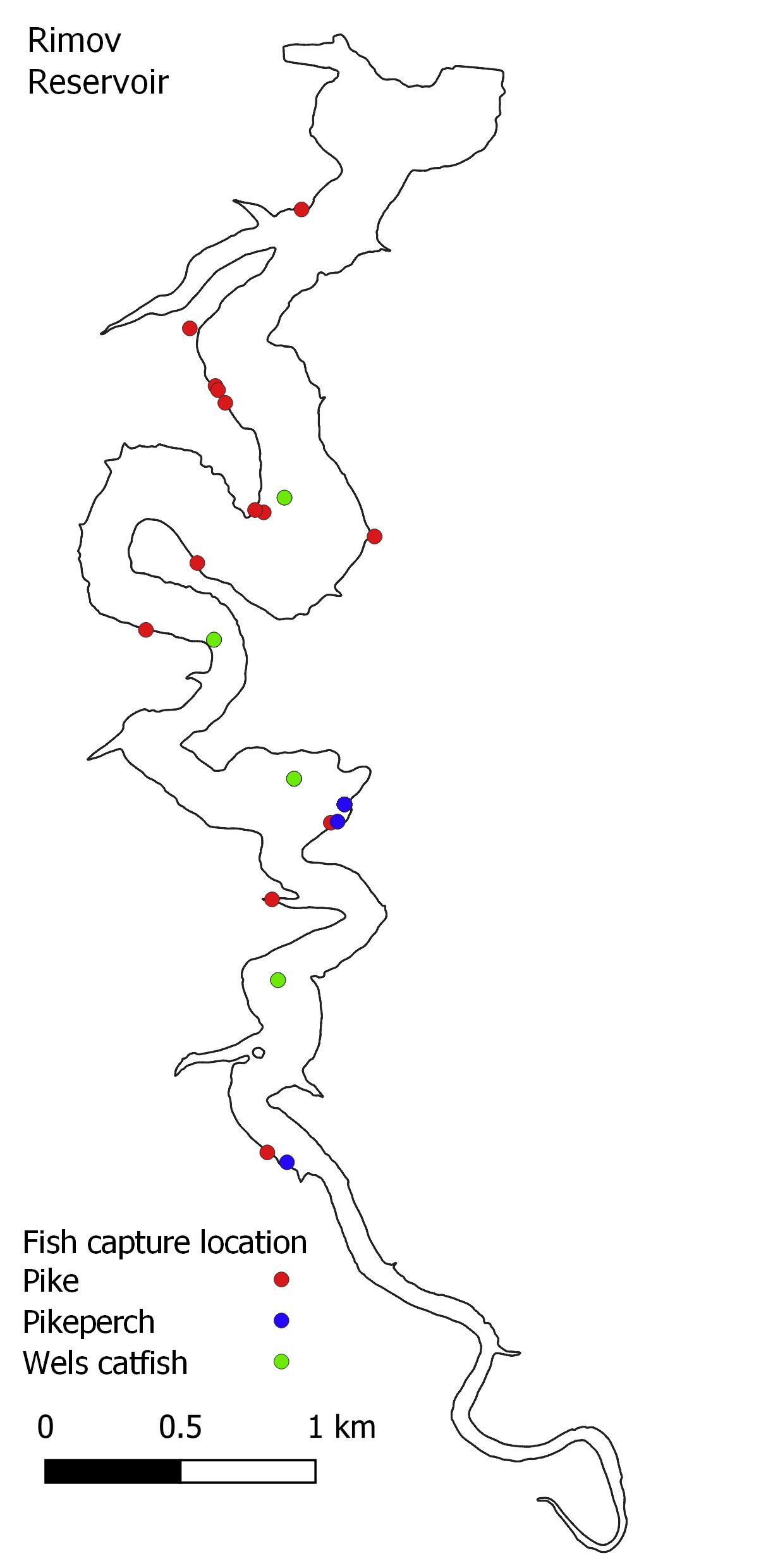

**Figure S2.** Predicted probabilities of reservoir section by season

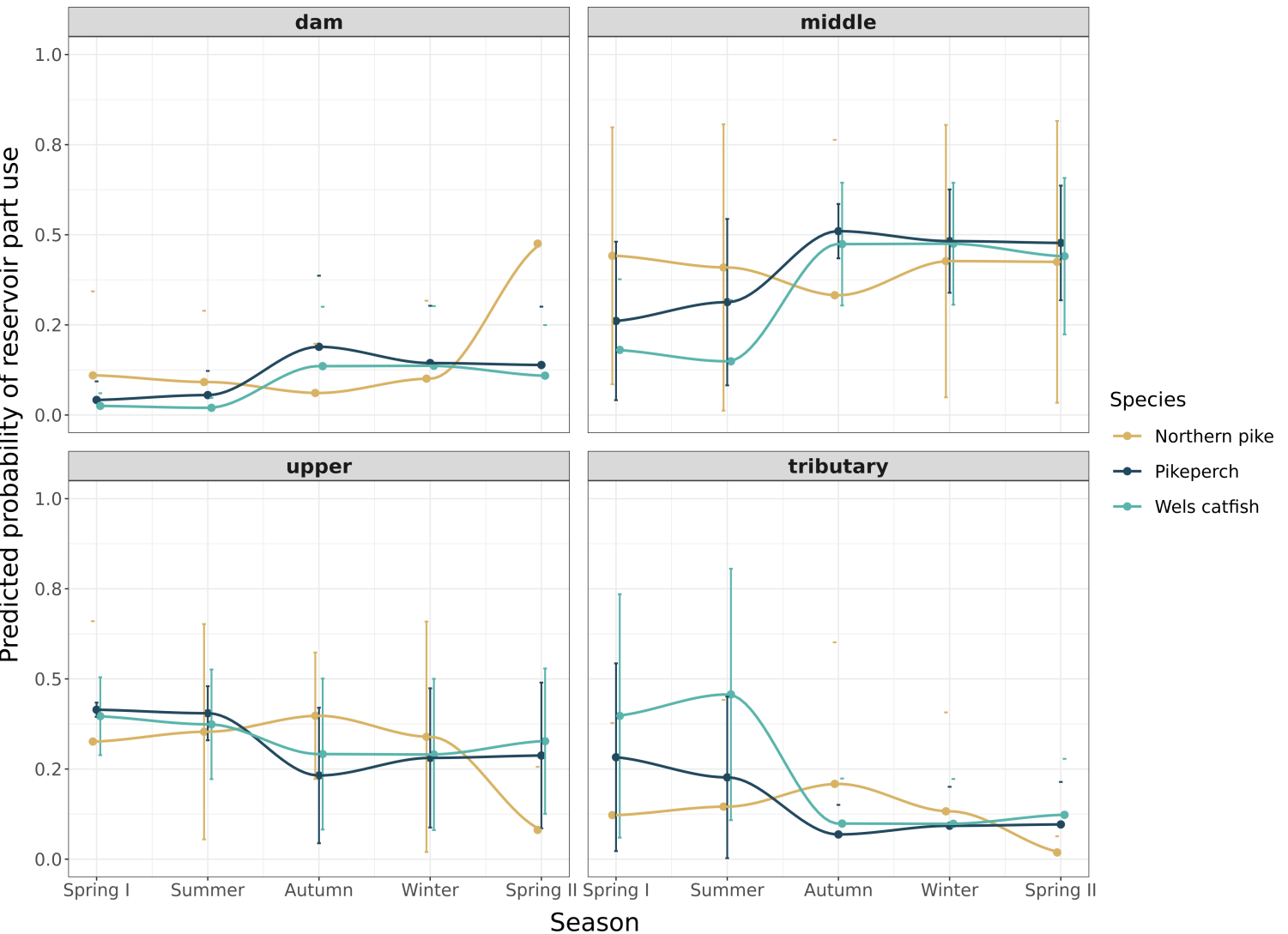

**Figure S3.** Predicted probabilities of reservoir section use as a function of temperature

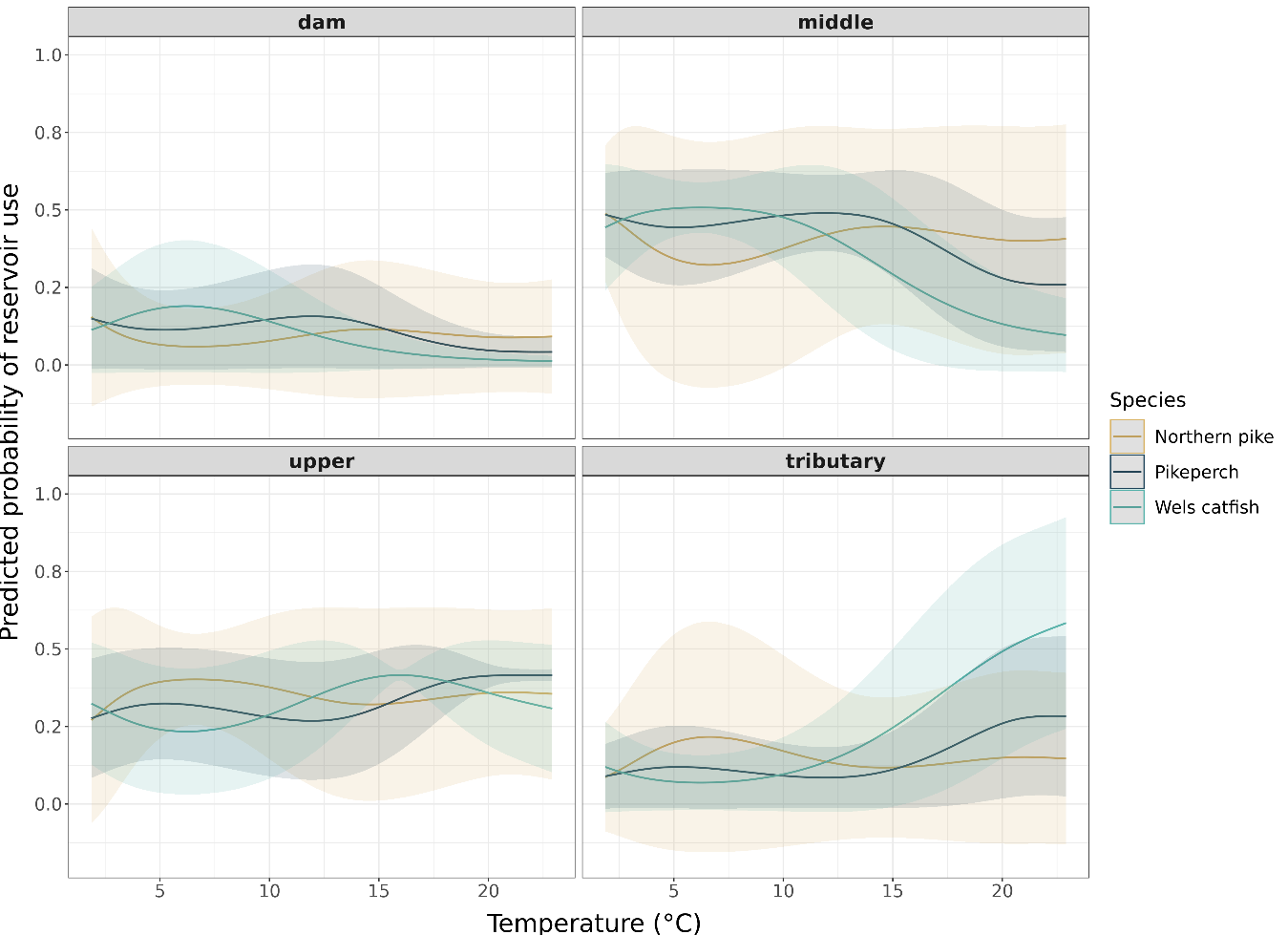

**Figure S4.** Predicted probabilities of depth use by season

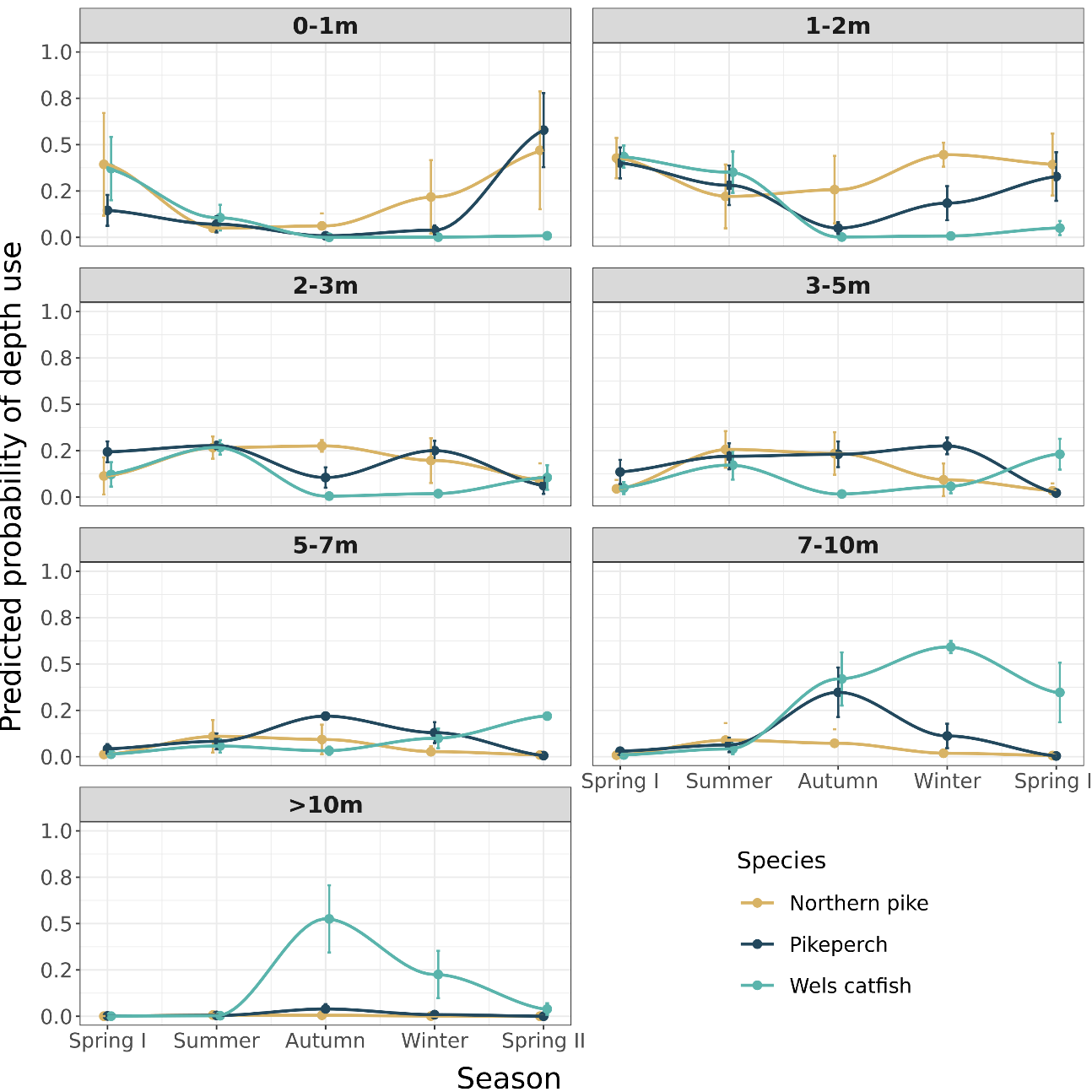
